# The mouse *Cdh23^ahl^* allele drives early cochlear dysfunction and phenotypic heterogeneity before progressive hearing loss

**DOI:** 10.64898/2026.11.02.763796

**Authors:** Sherylanne Newton, Marisa Flook, Christopher Shults, Pavandeep Gill, Carlos Aguilar, Wei Song, Michelle Stewart, Sara Wells, Ronna Hertzano, Jing-Yi Jeng, Michael R Bowl

**Affiliations:** UCL Ear Institute, University College London, 332 Gray’s Inn Road, London, UK; Section on Omics and Translational Science of Hearing, National Institute on Deafness and Other Communication Disorders, Bethesda, Maryland, USA; Mary Lyon Centre at MRC Harwell, Harwell Campus, UK; School of Biosciences, University of Sheffield, Sheffield, UK

**Keywords:** Co-isogenic mouse strains, C57BL/6N, Cadherin related 23, Cochlear transcriptomics, Hearing Loss, Phenotype expressivity, Mechanoelectrical transduction, Hair cell electrophysiology

## Abstract

The *Cdh23^ahl^* allele is present in many commonly used inbred mouse strains and is known to cause progressive hearing loss. However, increasing evidence suggests *Cdh23^ahl^*can also modify the severity and expressivity of other genetic phenotypes, extending its biological effects beyond the cochlea. Here, we define the phenotypic, physiological, and molecular consequences associated with *Cdh23^ahl^*by comparing C57BL/6N mice with a co-isogenic strain in which the allele was repaired by genome editing. We find the auditory phenotype begins earlier and is more variable than generally appreciated, with significant threshold elevations at 36–42 kHz already present by 1-month of age and progressive involvement of lower frequencies thereafter. Before these threshold elevations, inner hair cells at postnatal day 14 (P14) exhibit tonotopically distinct alterations in mechanoelectrical transduction, with reduced maximal transduction currents apically, but increased currents basally, while resting open probability and adaptation kinetics remain preserved. Cochlear transcription is also altered shortly after hearing onset: by P16, *Cdh23^ahl^*and repaired cochleae occupy distinct transcriptional states. By P30, transcriptional differences have expanded substantially, and *Cdh23^ahl^*cochleae show significantly increased inter-individual transcriptomic dispersion coincident with the emergence of variable high-frequency auditory dysfunction. These findings demonstrate that *Cdh23^ahl^* is not physiologically benign in young mice, but is associated with an altered cochlear state that precedes hearing loss. The early and variable consequences of this common allele provide a potential framework for understanding its effects on phenotype expressivity and have important implications for the design and interpretation of studies using mouse strains that carry *Cdh23^ahl^*.

**Summary:** The *Cdh23^ahl^* allele, common to several mouse strains, is known to cause progressive hearing loss, but it can also modify other genetic phenotypes. Using C57BL/6N mice, and a co-isogenic strain in which *Cdh23^ahl^* was repaired, the biological effects of *Cdh23^ahl^* were investigated. Results show the *Cdh23^ahl^* allele causes tonotopically distinct alterations in inner hair cell mechanotransduction and progressive changes in the cochlear transcriptome, leading to a variable auditory phenotype. The findings demonstrate *Cdh23^ahl^* is not a simple loss-of-function allele and provide insight into its ability to modify phenotype severity and expressivity, which has broad implications for studies employing *Cdh23^ahl^*-carrying strains.

## Introduction

Cadherin-23 (CDH23), also known as otocadherin, is an atypical member of the cadherin superfamily. Within stereocilia bundles located on the apical surface of inner ear sensory hair cells, CDH23 together with Protocadherin-15 (PCDH15) form filamentous tip-links that connect the top of one stereocilium to the side of a taller adjacent stereocilium^1–3^. In the cochlea, sound-induced movement of the bundles cause tip-links to stretch, opening mechanically gated mechanoelectrical transducer (MET) channels present near the lower end of the link^4–6^. This allows an influx of ions, leading to depolarization of the hair cell, which in turn leads to release of glutamate to elicit a neural impulse that travels to the brain^7^. As such, CDH23 is an essential component of the MET apparatus, permitting sound to be converted into electrical signals that enable hearing. Consistent with this essential function, pathogenic variants in *CDH23* cause severe congenital hearing impairment in humans and mice^8–11^. However, more than 20 inbred mouse strains carry a fixed *Cdh23* allele (*Cdh23^ahl^*), including C57BL/6J and C57BL/6N^12^, and intriguingly these mice exhibit relatively normal auditory thresholds early in life before developing a progressive hearing loss that begins in the higher frequency range^12–15^. This apparent disconnect between the fundamental role of CDH23 in mechanotransduction and the delayed auditory phenotype associated with *Cdh23^ahl^* remains incompletely understood.

The *Cdh23^ahl^* allele results in incomplete skipping of exon 9 and has long been recognized as a major contributor to age-related hearing loss in the mouse strains that carry it^13^. Progressive auditory dysfunction is accompanied by multiple cochlear pathologies, including outer hair cell loss, degeneration of the spiral ligament and loss of spiral ganglion neurons^16^. Given that substantial threshold elevations have generally been associated with later ages, the *Cdh23^ahl^*allele is frequently treated as inconsequential in studies performed in young C57BL/6-derived mice.

However, increasing evidence challenges this assumption, such as studies showing that the *Cdh23^ahl^* allele can interact with other genetic variants to alter phenotype severity and expressivity. The allele potentiates hearing loss associated with mutations in *Nptn*, *Sod1*, *Atp2b2*, *Ocm,* and *Wbp2*^17–21^. We previously found an even broader interaction with loss of *Embigin*: *Embigin*-null mice maintained on the C57BL/6N.*Cdh23^ahl^* background developed hearing loss together with cardiac and brain abnormalities and reduced viability, whereas these phenotypes were absent or substantially ameliorated when the same mutation was maintained on a C57BL/6N background in which *Cdh23^ahl^* had been repaired^22^. These observations suggest the *Cdh23^ahl^* allele can influence biological phenotypes earlier, and potentially beyond, its recognized contribution to age-related hearing loss.

Understanding these effects is particularly important because C57BL/6-derived strains are among the most extensively used genetic backgrounds in biomedical research. C57BL/6N, for example, is the background strain used by the International Mouse Phenotyping Consortium for the many thousands of genetically modified (knockout) mouse lines they have generated^23,24^. Thus, biological effects of the *Cdh23^ahl^* allele that precede overt hearing loss could influence the penetrance or expressivity of experimentally introduced mutations and contribute to reproducibility differences between models or laboratories.

A powerful approach to defining these effects is provided by a co-isogenic C57BL/6N strain in which we previously corrected the *Cdh23^ahl^* allele by CRISPR/Cas9-mediated homology-directed repair^14^. The resulting C57BL/6N.*Cdh23^c.753A>G^*strain (hereafter referred to as B6N.*Cdh23^repair^*) differs from the parental C57BL/6N strain (hereafter referred to as B6N.*Cdh23^ahl^*) at the repaired locus while retaining the same genetic background, enabling the consequences associated with *Cdh23^ahl^* to be examined without any confounding effects from background genetic variation.

Here, we used these co-isogenic strains to determine when and how the *Cdh23^ahl^* allele begins to alter biological function. We combined broad phenotyping and longitudinal auditory measurements with direct recordings of hair-cell MET currents and cochlear transcriptomic profiling. We find the effects of the *Cdh23^ahl^* allele emerge substantially earlier than generally appreciated. High-frequency auditory thresholds become abnormal and highly variable in young mice. Prior to this, shortly after hearing onset, inner hair cells exhibit tonotopically distinct changes in MET current magnitude and cochlear gene-expression programs are altered. By 1-month of age, B6N.*Cdh23^ahl^*cochleae also display striking inter-individual transcriptomic heterogeneity that parallels the variability observed in auditory phenotype. Together, these findings identify *Cdh23^ahl^* as a biologically active modifier of cochlear state before substantial hearing loss develops and provide a framework for understanding how this common background allele can influence phenotype expressivity in genetically modified mice.

## Results

### *Cdh23^ahl^* causes early-onset and highly variable high-frequency hearing loss

The *Cdh23^ahl^* allele is well established as a major contributor to progressive age-related hearing loss in C57BL/6-derived mouse strains, with high-frequency threshold elevations generally becoming prominent during the first several months of life^12^. Given that CDH23 is required for hair-cell mechanotransduction from the onset of hearing (∼P12), we asked whether the auditory consequences of *Cdh23^ahl^*might emerge earlier than previously reported^25^.

To define the trajectory of hearing loss associated with *Cdh23^ahl^*, we compiled auditory brainstem response (ABR) measurements collected from B6N.*Cdh23^ahl^*mice and co-isogenic B6N.*Cdh23^repair^* mice across multiple studies performed in our laboratory between 2016 and 2023. ABR thresholds were available for 8, 16, and 32 kHz pure-tone stimuli at 1, 3, 6, 9, 12, 15, and/or 18 months of age. The combined dataset included 170 B6N.*Cdh23^ahl^* mice (86 male, 84 female), of which 95 were assessed at more than one time point, and 90 B6N.*Cdh23^repair^*mice (42 male, 48 female), of which 84 were assessed longitudinally.

Because sex differences have been reported in age-related hearing loss in mice, male and female animals were initially analyzed separately. Auditory trajectories were broadly similar between sexes, with only a modest difference in 32-kHz thresholds at 3 months of age [female median = 50 dB SPL, IQR = 25 dB SPL; male median = 60 dB SPL, IQR = 25 dB SPL; *P* = 0.0372, Mann–Whitney test with Holm–Sidak correction (Supplementary Figure 1)]. We therefore found no evidence for a consistent sex-dependent difference in the onset or progression of the *Cdh23^ahl^*-associated auditory phenotype.

The dataset revealed the expected progressive high-frequency hearing loss in B6N.*Cdh23^ahl^* mice and preservation of high-frequency hearing in B6N.*Cdh23^repair^* mice (Figure 1A,B). Unexpectedly, however, differences were evident considerably earlier than anticipated. At one month of age, 32 kHz thresholds were already more variable among B6N.*Cdh23^ahl^* mice than among B6N.*Cdh23^repair^* mice (IQR = 15 versus 10 dB SPL), with a subset of B6N.*Cdh23^ahl^* animals exhibiting elevated thresholds. By three months, this divergence was striking: 32 kHz thresholds in B6N.*Cdh23^ahl^* mice ranged from 20 dB SPL to no detectable response at the maximum 90 dB SPL stimulus (IQR = 25 dB SPL), whereas thresholds in B6N.*Cdh23^repair^* mice remained tightly distributed between 15 and 45 dB SPL (IQR = 5 dB SPL; Figure 1A,B).

**Figure 1.**
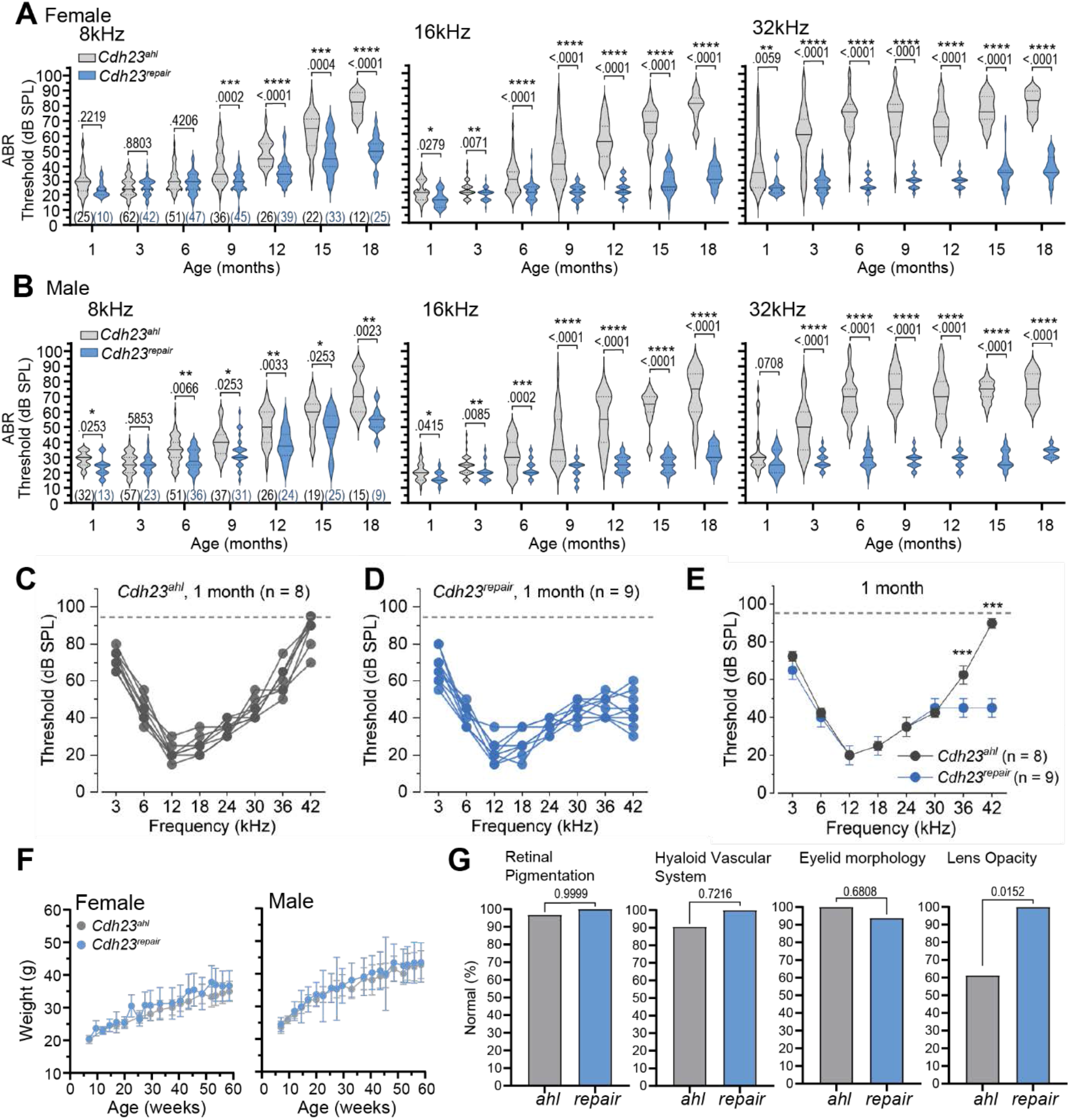
*Cdh23^ahl^*causes with early-onset and increased variability of progressive high-frequency hearing loss. **(A,B)** Auditory brainstem response (ABR) thresholds measured in female **(A)** and male **(B)** B6N.*Cdh23^ahl^* and B6N.*Cdh23^repair^* mice in response to 8, 16, and 32 kHz pure-tone stimuli between 1 and 18 months of age. Data were compiled from multiple cohorts assessed in our laboratory between 2016 and 2023. Violin plots show the distribution of individual thresholds at each age. Numbers of mice included at each time point are shown below the corresponding 8 kHz distributions. Between-genotype comparisons were performed using Mann–Whitney tests with Holm–Šídák correction for multiple comparisons; adjusted *P* values are shown above the corresponding distributions. The dashed line indicates the maximum stimulus level of the ABR system. B6N.*Cdh23^ahl^* mice show early and progressively elevated high-frequency thresholds, together with a broad distribution of 32-kHz thresholds at younger ages. **(C,D)** Individual ABR thresholds from one-month-old B6N.*Cdh23^ahl^* **(C)** and B6N.*Cdh23^repair^* **(D)** mice measured across an extended frequency range of 3–42 kHz. **(E)** Comparison of ABR thresholds from the animals shown in **(C,D)**. Data are presented as median ± median absolute deviation (MAD). At one month of age, thresholds were comparable between genotypes from 3 to 30 kHz but were significantly elevated in B6N.*Cdh23^ahl^* mice at 36 and 42 kHz. Statistical comparisons were performed using Kruskal–Wallis ANOVA. Data include five one-month-old animals of each genotype reported previously^26^. Dashed lines indicate the maximum stimulus level of the ABR system. **(F)** Longitudinal body-weight measurements from male and female B6N.*Cdh23^ahl^* and B6N.*Cdh23^repair^* mice from 7 to 59 weeks of age. No significant genotype-dependent differences in growth trajectory were detected. **(G)** Incidence of lens opacity in B6N.*Cdh23^ahl^* (n = 31) and B6N.*Cdh23^repair^* (n = 16) mice assessed at 58 weeks of age. Lens opacity affecting one or both eyes was observed in approximately 40% of B6N.*Cdh23^ahl^* mice and was not detected in B6N.*Cdh23^repair^* mice. Statistical comparison was performed using a two-sided Fisher’s exact test with correction for multiple comparisons. \**P* < 0.05, \*\**P* < 0.01, \*\*\**P* < 0.001, \*\*\*\**P* < 0.0001.

To further characterise the apparent genotype-associated differences in inter-animal variability, differences in threshold variance between genotypes were assessed. Together, these analyses characterised the persistence of inter-animal variability in hearing loss progression (Supplementary Table 1). At 32 kHz, B6N.*Cdh23^ahl^* mice continued to show broad threshold distributions at 6 (Brown–Forsythe test: F = 841.54, P = 3.24×10^−63^), 9 (Brown–Forsythe test: F= 819.81, P = 7.88×10^−48^), and 12 (Brown–Forsythe test: F = 436.96, P = 5.59×10^−30^) months of age. By 15–18 months, the distribution narrowed as an increasing proportion of animals reached severe-to-profound threshold elevations or failed to respond at the maximum stimulus level. The variability at younger ages was not explained by litter or postnatal cage density: 32 kHz thresholds at three months showed no significant relationship with litter/cage density between P12 and weaning (*R*² = 0.0234, *P* = 0.097; Supplementary Figure 2).

The auditory phenotype progressed from high to lower frequencies with age. At 16 kHz, thresholds were similar between strains at one and three months, but became elevated and more heterogeneous in B6N.*Cdh23^ahl^*mice from six months onward (Brown–Forsythe test: F = 47.32, P = 1.23×10^−10^) (Figure 1A,B). At 8 kHz, thresholds were likewise comparable early in life and increased in both strains beginning at approximately six months, although elevations were generally greater in B6N.*Cdh23^ahl^* mice. Thus, repair of *Cdh23^ahl^* largely prevented progressive high- and mid-frequency hearing loss, but did not prevent the late-onset low-frequency threshold elevation, indicating that the latter is not solely attributable to *Cdh23^ahl^*.

To more precisely define the earliest auditory phenotype, we next assessed an independent cohort of one-month-old mice using an extended range of pure-tone stimuli from 3 to 42 kHz. Because the longitudinal dataset showed no consistent sex-dependent effect, male and female mice were combined for this analysis. Thresholds from 3 to 30 kHz were similar between strains. In contrast, B6N.*Cdh23^ahl^* mice already exhibited substantially elevated thresholds at 36 kHz (62.5 ± 5 versus 45 ± 5 dB SPL) and 42 kHz (90 ± 2.5 versus 45 ± 5 dB SPL; both *P* < 0.001; Figure 1C–E).

Together, these findings demonstrate that the auditory consequences of *Cdh23^ahl^* begin earlier than generally appreciated and are initially restricted to the extreme high-frequency region of the cochlea. Importantly, the phenotype is characterized not only by progressive threshold elevation, but also by pronounced inter-individual variability in its onset and severity. Repair of *Cdh23^ahl^*largely eliminates both features, identifying the allele as a major driver of early high-frequency auditory dysfunction and phenotypic heterogeneity.

### *Cdh23^ahl^* is associated with increased lens opacity in aged mice

Previous studies have demonstrated that *Cdh23^ahl^* can modify phenotypes outside the auditory system, as such we asked whether repair of the allele was associated with broader phenotypic differences during aging. We therefore performed standardized phenotyping of 41 B6N.*Cdh23^ahl^* mice (20 female, 21 male) and 27 B6N.*Cdh23^repair^*mice (14 female, 13 male), including longitudinal body weight measurements and assessments of neurobehavioral, cardiovascular, metabolic, hematologic, clinical chemistry, body-composition, and ocular phenotypes (Supplementary File 1).

The two strains showed similar growth trajectories from 7 to 59 weeks of age (Figure 1F), and no major differences were detected in cardiac or neurobehavioral phenotypes. Several statistically significant differences were observed in clinical chemistry and hematologic measures; however, these were generally modest in magnitude. These are detailed in Supplementary File 1 and Supplementary Figures 3 and 4.

The most prominent non-auditory phenotype was an increased incidence of lens opacity. At 58 weeks of age, lens opacity in one or both eyes was observed in 38.7% of assessed B6N.*Cdh23^ahl^* mice, but in none of the B6N.*Cdh23^repair^* mice (*P* = 0.0038, Fisher’s exact test; Figure 1G). Given that ocular morphology was assessed only at this age, the time of onset of the phenotype could not be determined.

### *Cdh23^ahl^* alters inner hair cell mechanoelectrical transduction in a tonotopically distinct manner

Given that CDH23 is an essential component of stereociliary tip links, we asked whether *Cdh23^ahl^* affects mechanoelectrical transduction (MET) in inner hair cells (IHCs) before hearing loss develops. For this, we performed whole-cell patch-clamp recordings from IHCs at P14, shortly after the onset of hearing and before the high-frequency threshold elevations can be detected at one month of age.

To determine whether any effects differed along the cochlear tonotopic axis, MET currents were recorded from IHCs in both apical (9–12 kHz) and basal (40–45 kHz) regions of B6N.*Cdh23^ahl^* and B6N.*Cdh23^repair^*cochleae. Post-hearing MET recordings are generally restricted to more apical regions because of technical challenges; however, here for the first time we were able to elicit and record MET currents from both apical and basal IHCs using a fluid-jet to elicit transducer currents (Figure 2A–D).

**Figure 2.**
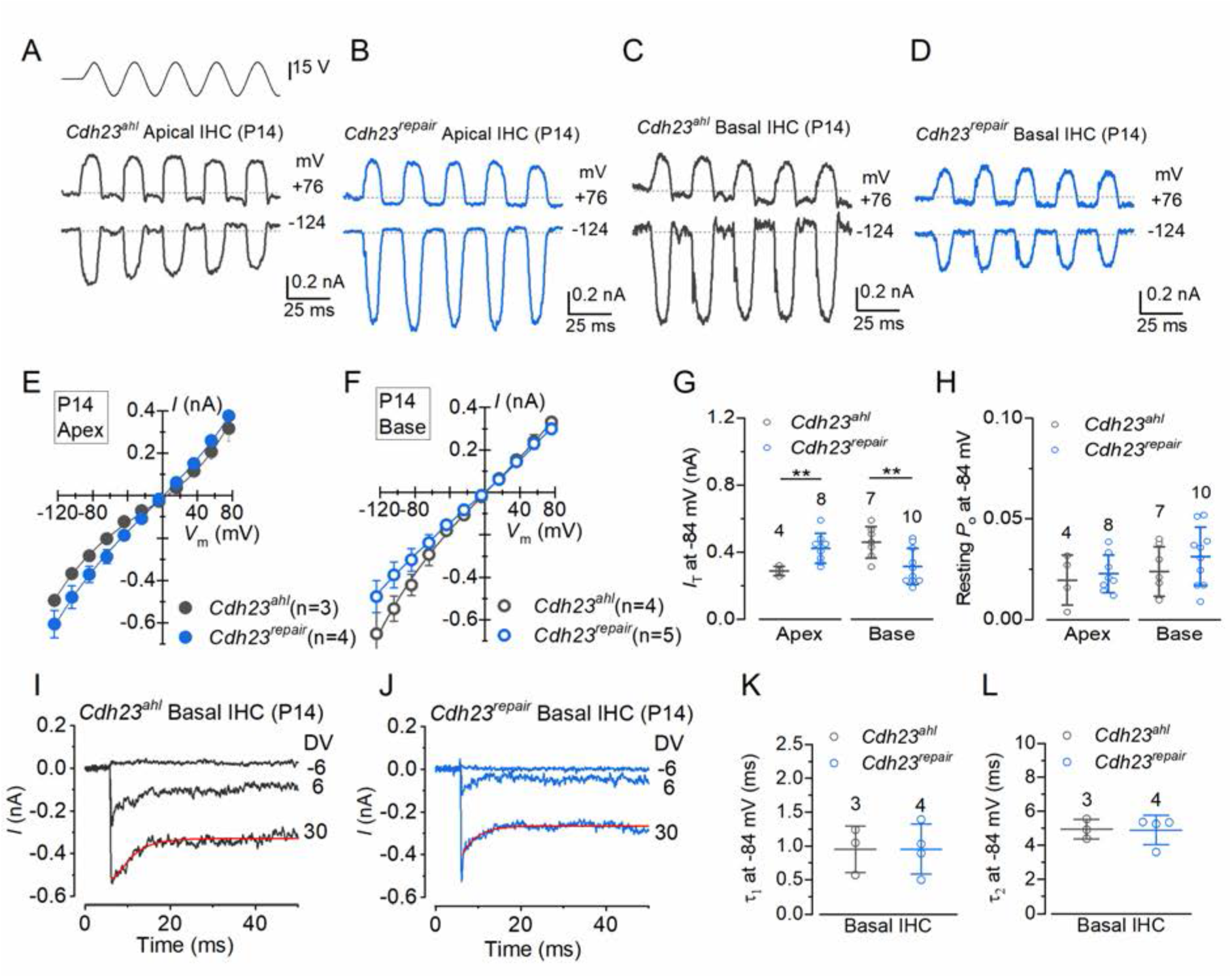
*Cdh23^ahl^* alters inner hair cell mechanoelectrical transduction in a tonotopically distinct manner. Mechanoelectrical transduction (MET) currents were recorded from apical (9-12 kHz) and basal (40-45 kHz) inner hair cells (IHCs) of P14 B6N.*Cdh23^ahl^* and B6N.*Cdh23^repair^* mice using whole-cell patch clamp and fluid-jet stimulation. **(A–D)** Representative MET currents elicited by a 50-Hz sinusoidal fluid-jet stimulus in apical **(A,B)** and basal **(C,D)** from a holding potential of −84 mV in IHCs. Positive driver voltages displaced the hair bundle in the excitatory direction and negative driver voltages in the inhibitory direction. Representative currents are shown at holding potentials of +76 and −124 mV. Dashed lines indicate the holding current. **(E,F)** Current–voltage relationships for maximal MET currents recorded from apical **(E)** and basal **(F)** IHCs over membrane potentials from −124 to +76 mV. Data are shown as mean ± SEM and fitted using a nonlinear current–voltage function^27^. MET currents reversed close to 0 mV in both genotypes and cochlear regions. **(G)** Maximal MET-current amplitude measured at −84 mV. B6N.*Cdh23^ahl^*IHCs exhibited significantly smaller maximal MET currents than B6N.*Cdh23^repair^*IHCs in the apical cochlea (*P* = 0.015), but significantly larger maximal currents in the basal cochlea (*P* = 0.012). **(H)** Resting open probability (Po) of the MET current at −84 mV. No significant genotype-dependent differences were detected in either apical (*P* = 0.636) or basal (*P* = 0.289) IHCs. **(I,J)** Representative MET currents from basal IHCs elicited at −84 mV using force-step stimuli. Positive fluid-jet driver voltages (DV) of 2 mV steps elicited inward currents that adapted during sustained stimulation. Maximum current responses were recorded at a DV of 30 mV, fitted with a Boltzmann function to derive adaptation time constants (red lines). **(K,L)** Fast (τ1) **(K)** and slow (τ2) **(L)** decay time constants of MET-current adaptation in basal IHCs. No significant differences were detected between B6N.*Cdh23^ahl^* and B6N.*Cdh23^repair^* IHCs (τ1, *P* = 0.996; τ2, *P* = 0.917). Numbers of recorded cells are indicated above the data, and each cell was recorded from a different animal. Unless otherwise indicated, data are presented as mean ± SD.

Saturating MET currents were first elicited using a 50-Hz sinusoidal fluid-jet stimulus to displace the hair bundle in the excitatory and inhibitory directions. Current–voltage relationships were measured over membrane potentials ranging from −124 to +76 mV. MET currents reversed close to 0 mV in both strains and cochlear regions [apical IHCs: B6N.*Cdh23^ahl^*, 3.2 ± 1.3 mV; B6N.*Cdh23^repair^*, 2.3 ± 1.4 mV; basal IHCs: B6N.*Cdh23^ahl^*, 2.0 ± 1.7 mV; B6N.*Cdh23^repair^*, 1.2 ± 1.8 mV (Figure 2E,F)], consistent with the expected non-selective cation conductance of the MET channel.

In contrast, the maximal MET current measured at −84 mV differed significantly between genotypes, and the direction of this effect depended on cochlear location. In apical IHCs, maximal MET currents were smaller in B6N.*Cdh23^ahl^*than in B6N.*Cdh23^repair^* mice (*P* = 0.015). Strikingly, the opposite relationship was observed in basal IHCs, where maximal MET currents were larger in B6N.*Cdh23^ahl^* than in B6N.*Cdh23^repair^*mice (*P* = 0.012; Figure 2G). Thus, *Cdh23^ahl^* does not produce a uniform reduction in MET current, but instead alters current magnitude in a tonotopically distinct manner.

We next asked whether *Cdh23^ahl^* altered other properties of the MET response. Resting open probability measured at −84 mV did not differ significantly between genotypes in either apical or basal IHCs (apical, *P* = 0.636; basal, *P* = 0.289; Figure 2H). Similarly, analysis of MET-current adaptation in basal IHCs using force-step stimuli revealed no significant differences in either the fast (τ1) or slow (τ2) decay time constants between B6N.*Cdh23^ahl^* and B6N.*Cdh23^repair^* mice (τ1, *P* = 0.996; τ2, *P* = 0.917; Figure 2I–L).

Together, these findings demonstrate that *Cdh23^ahl^* alters the magnitude of IHC MET currents shortly after hearing onset without abolishing mechanotransduction or detectably altering resting channel open probability or adaptation kinetics. Notably, the effect differs along the cochlear tonotopic axis, with increased maximal MET currents in basal IHCs of B6N.*Cdh23^ahl^* mice, corresponding to the cochlear region in which auditory dysfunction first emerges.

### *Cdh23^ahl^* alters the cochlear transcriptome shortly after hearing onset and is associated with increasing inter-individual heterogeneity

The presence of altered MET currents at P14 suggested that *Cdh23^ahl^* affects cochlear biology before substantial auditory threshold elevations are apparent. To determine whether these early physiological changes are accompanied by alterations in cochlear gene expression, we performed bulk RNA sequencing of whole cochleae, with the vestibular organs removed, from B6N.*Cdh23^ahl^* and B6N.*Cdh23^repair^* mice at P16 and P30. These time points were selected to capture molecular changes shortly after hearing onset (P16) and at an age when the earliest high-frequency auditory deficits are emerging (P30). At P16, seven B6N.*Cdh23^ahl^* samples (4 male, 3 female) were compared with seven B6N.*Cdh23^repair^* samples (3 male, 4 female); at P30, seven B6N.*Cdh23^ahl^* samples (3 male, 4 female) were compared with six B6N.*Cdh23^repair^* samples (4 male, 2 female).

The *Cdh23^ahl^* allele has been reported to cause incomplete in-frame skipping of exon 9 (ENSMUSE00001269485). Visualization of RNA-seq reads from the P16 samples using the Integrative Genomics Viewer confirmed incomplete exon 9 skipping in B6N.*Cdh23^ahl^* cochleae, whereas exon 9 was retained in transcripts from B6N.*Cdh23^repair^* cochleae (Figure 3A), confirming the expected molecular consequence of allele repair.

**Figure 3.**
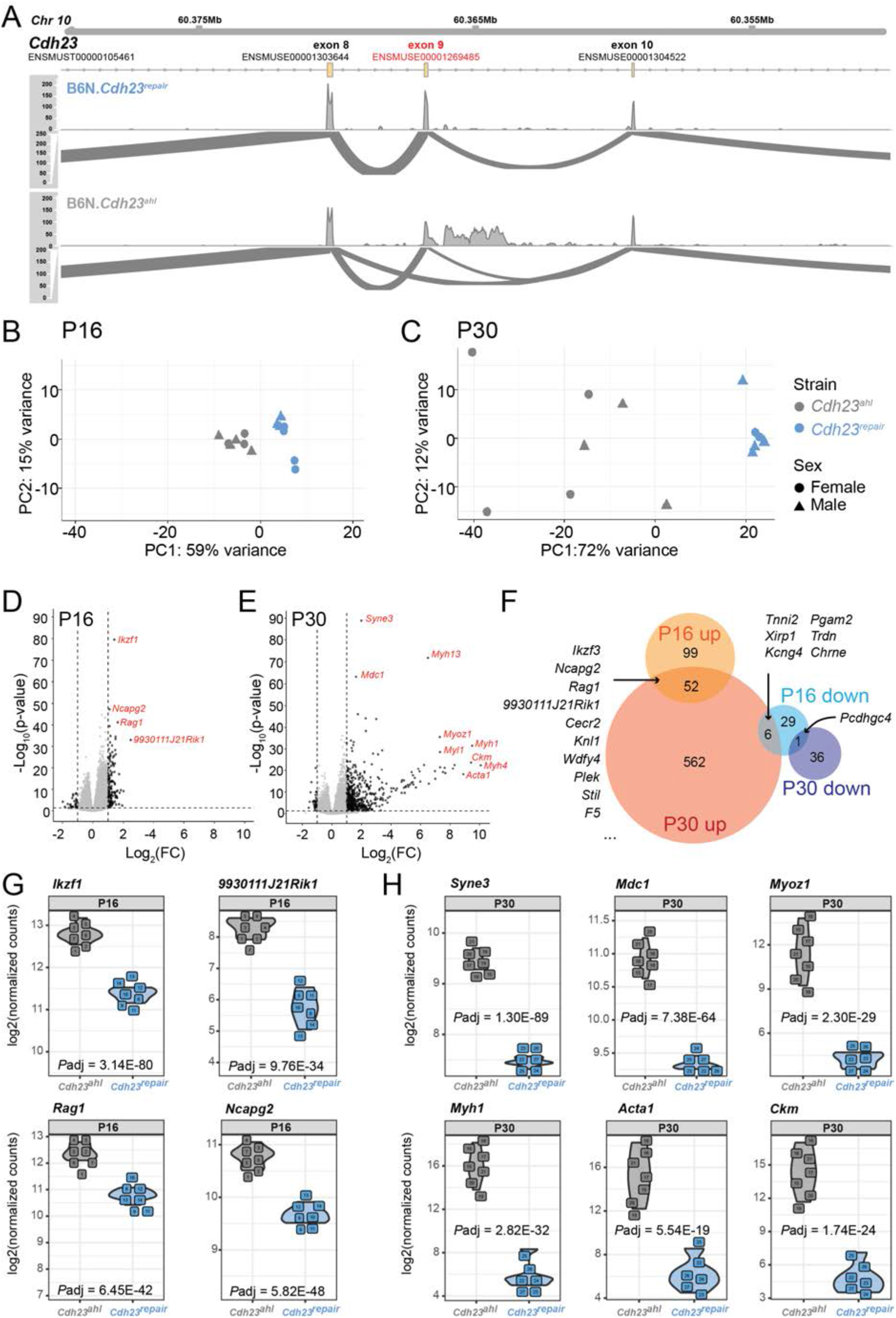
*Cdh23^ahl^*-associated transcriptional changes emerge shortly after hearing onset and expand by P30. **(A)** RNA-sequencing reads spanning the *Cdh23* locus in P16 B6N.*Cdh23^ahl^* (n = 7) and B6N.*Cdh23^repair^* (n = 7) cochleae, visualized using the Integrative Genomics Viewer (IGV) after merging reads by genotype. Arcs represent reads spanning exon junctions, with arc height proportional to read number; peaks indicate sequencing coverage. B6N.*Cdh23^ahl^* samples show incomplete skipping of exon 9, whereas exon 9 is retained in B6N.*Cdh23^repair^* transcripts. **(B,C)** Principal component analysis of bulk cochlear RNA-seq samples at P16 **(B)** and P30 **(C)** using the 500 most variable genes. At P16, samples separate according to genotype. At P30, B6N.*Cdh23^repair^* samples remain closely grouped, whereas B6N.*Cdh23^ahl^* samples show substantially greater inter-individual dispersion. **(D,E)** Volcano plots showing differential gene expression between B6N.*Cdh23^ahl^* and B6N.*Cdh23^repair^*cochleae at P16 **(D)** and P30 **(E)**. Differentially expressed genes (DEGs) were defined as |log2 fold change| >1 and adjusted *P* <0.05. At P16, 187 genes were differentially expressed (151 upregulated, 36 downregulated); at P30, 657 genes were differentially expressed (620 upregulated, 37 downregulated). Selected highly significant DEGs are labeled. **(F)** Euler diagram showing overlap among genes up- or downregulated in B6N.*Cdh23^ahl^*relative to B6N.*Cdh23^repair^* cochleae at P16 and P30. Fifty-two genes were upregulated and one gene was downregulated at both ages; six genes downregulated at P16 were upregulated at P30. **(G,H)** Normalized read counts for representative DEGs at P16 **(G)** and P30 **(H)**. Individual B6N.*Cdh23^ahl^* and B6N.*Cdh23^repair^*samples show relatively consistent expression at P16, whereas greater inter-individual variability is evident among B6N.*Cdh23^ahl^* samples at P30.

Principal component analysis (PCA) based on the 500 most variable genes revealed genotype-associated differences at both ages. At P16, B6N.*Cdh23^ahl^* and B6N.*Cdh23^repair^*samples formed distinct clusters, demonstrating that the cochlear transcriptome is already altered in *Cdh23^ahl^* mice shortly after hearing onset (Figure 3B). However, PERMDISP analysis revealed no significant difference in multivariate dispersion between genotypes at P16 (F = 3.2631, P = 0.1017), suggesting that the observed separation was not accompanied by a detectable difference in within-genotype transcriptomic variability. By P30, B6N.*Cdh23^repair^*samples remained tightly grouped, whereas B6N.*Cdh23^ahl^* samples were substantially more dispersed (Figure 3C). Consistently, PERMDISP analysis demonstrated significantly greater multivariate dispersion in B6N.Cdh23*^ahl^*samples than in B6N.Cdh23*^repair^* samples (F = 5.8714, P = 0.034) (Supplementary Figure 5) Thus, in addition to genotype-dependent differences in gene expression, the transcriptomes of individual B6N.*Cdh23^ahl^* cochleae become increasingly heterogeneous by one month of age.

Differential expression analysis further demonstrated an expansion of the transcriptional differences between genotypes from P16 to P30. At P16, 187 genes were differentially expressed in B6N.*Cdh23^ahl^* relative to B6N.*Cdh23^repair^* cochleae (|log2 fold change| >1.0, adjusted *P* <0.05), including 151 upregulated and 36 downregulated genes (Figure 3D; Supplementary File 2). By P30, 657 genes were differentially expressed, comprising 620 upregulated and 37 downregulated genes (Figure 3E; Supplementary File 2). Thus, the increase in transcriptional differences at P30 was driven predominantly by genes with increased expression in B6N.*Cdh23^ahl^*cochleae.

Most differentially expressed genes were specific to one developmental stage. Fifty-two genes were upregulated and one gene was downregulated in B6N.*Cdh23^ahl^* cochleae at both P16 and P30, whereas six genes that were downregulated at P16 were upregulated at P30 (Figure 3F; Supplementary File 2). The 53 genes showing concordant differential expression at both ages were not significantly enriched for a Gene Ontology biological-process category. STRING analysis, however, identified two small groups of functionally related proteins associated with cell-cycle processes (*Kif11, Tubb1, Brca1, Knl1, Stil, Mis18bp1, Ncapg2,* and *Kif14*; adjusted *P* = 1.1 × 10^−4) and regulation of immune-system processes (*Il7r, Il18rap, Sele, Rag1, Il1rl1, Ikzf3,* and *Tlr8*; adjusted *P* = 9.7 × 10^−4).

Examination of individual normalized read counts reinforced the age-dependent increase in inter-individual variability. At P16, expression of representative differentially expressed genes was relatively consistent among individual samples of both genotypes (Figure 3G). At P30, B6N.*Cdh23^repair^* samples remained comparatively consistent, whereas expression levels among individual B6N.*Cdh23^ahl^* samples varied substantially for representative differentially expressed genes (Figure 3H).

Together, these data demonstrate that *Cdh23^ahl^* is associated with cochlear transcriptional changes shortly after hearing onset, before substantial auditory dysfunction is apparent. Over the subsequent two weeks, these differences expand and are accompanied by increasing inter-individual transcriptomic heterogeneity. This molecular heterogeneity emerges during the same developmental interval in which variability in high-frequency auditory thresholds becomes apparent, suggesting that molecular and functional phenotypic variability develop in parallel in B6N.*Cdh23^ahl^* mice.

### *Cdh23^ahl^* is associated with an evolving and increasingly interconnected cochlear transcriptional response

To define the biological processes associated with the early transcriptional response to *Cdh23^ahl^*, we first examined functional relationships among the 187 DEGs identified at P16. Of these, 139 could be mapped to proteins in the STRING database; the remaining genes included pseudogenes, non-coding and predicted genes, and multiple immunoglobulin variable and constant region genes. Of the mapped proteins, 67 (48.2%) formed a single connected network containing 215 interactions, substantially more than the 36 expected for a randomly selected protein set of this size (*P* < 1 × 10^−16^; Figure 4A).

**Figure 4.**
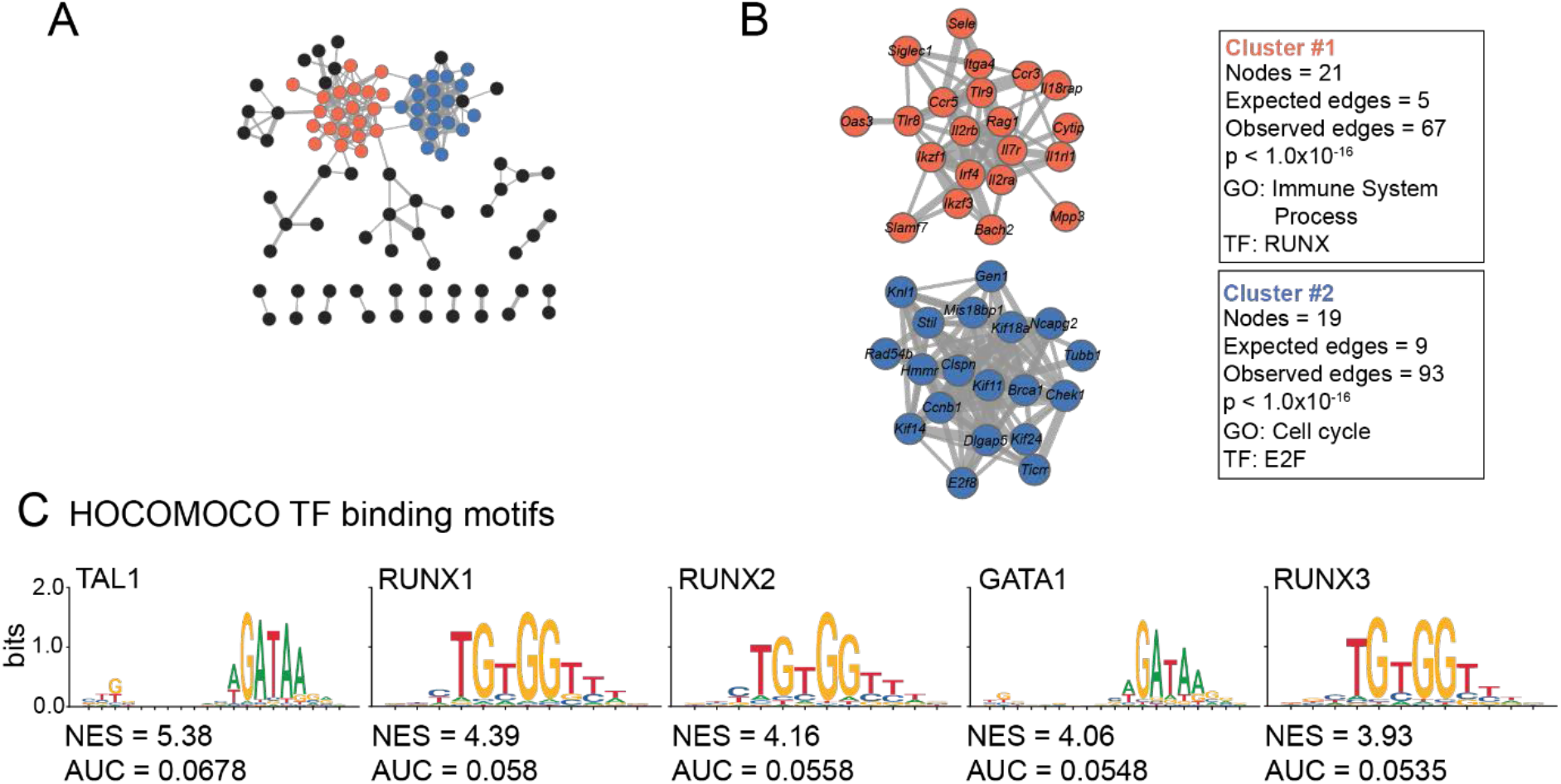
*Cdh23^ahl^*-associated genes form interconnected immune- and cell-cycle-related networks at P16. **(A)** STRING protein-interaction analysis of differentially expressed genes (DEGs) identified between P16 B6N.*Cdh23^ahl^* and B6N.*Cdh23^repair^*cochleae. Of 187 DEGs, 139 mapped to proteins in STRING and 67 formed a connected network. Nodes represent proteins and edges represent known or predicted protein interactions; edge thickness indicates interaction confidence. **(B)** Markov Cluster Algorithm (MCL) analysis identified two major modules containing >10 proteins within the P16 network. Cluster 1 was enriched for immune-system processes and Cluster 2 for cell-cycle processes. The most highly enriched transcription factor (TF) binding-motif family identified by RcisTarget for each cluster is shown. Network visualization was performed using Cytoscape. **(C)** TF binding-motif enrichment across the complete set of 187 P16 DEGs using RcisTarget and the HOCOMOCO mouse motif database. The five most highly enriched motifs are shown. Motif enrichment was assessed within the region spanning 500 bp upstream to 100 bp downstream of the transcription start site. NES, normalized enrichment score; AUC, area under the curve. Complete motif-enrichment results are provided in Supplementary File 4.

First, we analysed the data from P16. Gene Ontology analysis of the complete P16 DEG set identified enrichment for immune system processes (48 genes; adjusted *P* = 3.18 × 10^−10^), regulation of immune system processes (31 genes; adjusted *P* = 6.3 × 10^−5^), and defense response (29 genes; adjusted *P* = 1.7 × 10^−4^).

Markov clustering of the STRING network further identified two major interconnected modules. Cluster 1 contained 21 proteins and was enriched for immune system processes (adjusted *P* = 1.10 × 10^−9^), whereas Cluster 2 contained 19 proteins and was strongly enriched for cell-cycle processes (adjusted *P* = 4.33 × 10^−13^; Figure 4B; Supplementary File 3). Thus, the transcriptional response associated with *Cdh23^ahl^*shortly after hearing onset is characterized by coordinated changes in immune-related and cell-cycle-associated gene programs.

To identify candidate regulatory programs associated with these transcriptional changes, we performed transcription-factor binding motif enrichment analysis using RcisTarget, examining sequences from 500 bp upstream to 100 bp downstream of transcription start sites. The immune-associated Cluster 1 was enriched for RUNX-family motifs (NES = 6.86), whereas the cell-cycle-associated Cluster 2 was enriched for E2F-family motifs (NES ≥ 5.4; Figure 4B). Analysis of the complete P16 DEG set similarly identified enrichment for RUNX1, RUNX2, and RUNX3 motifs (NES = 4.39, 4.16, and 3.93, respectively), together with motifs associated with TAL1 (NES = 5.38) and GATA1 (NES = 4.06; Figure 4C; Supplementary File 4). These findings identify candidate regulatory signatures associated with the early transcriptional response but do not establish direct transcription-factor occupancy or activity.

By P30, the transcriptional response associated with *Cdh23^ahl^* was substantially larger and more interconnected. Of the 657 P30 DEGs, 568 mapped to proteins in STRING, of which 414 (72.9%) formed a single interconnected network containing 2,741 interactions compared with 663 expected by chance (*P* < 1 × 10^−16^; Figure 5A).

**Figure 5.**
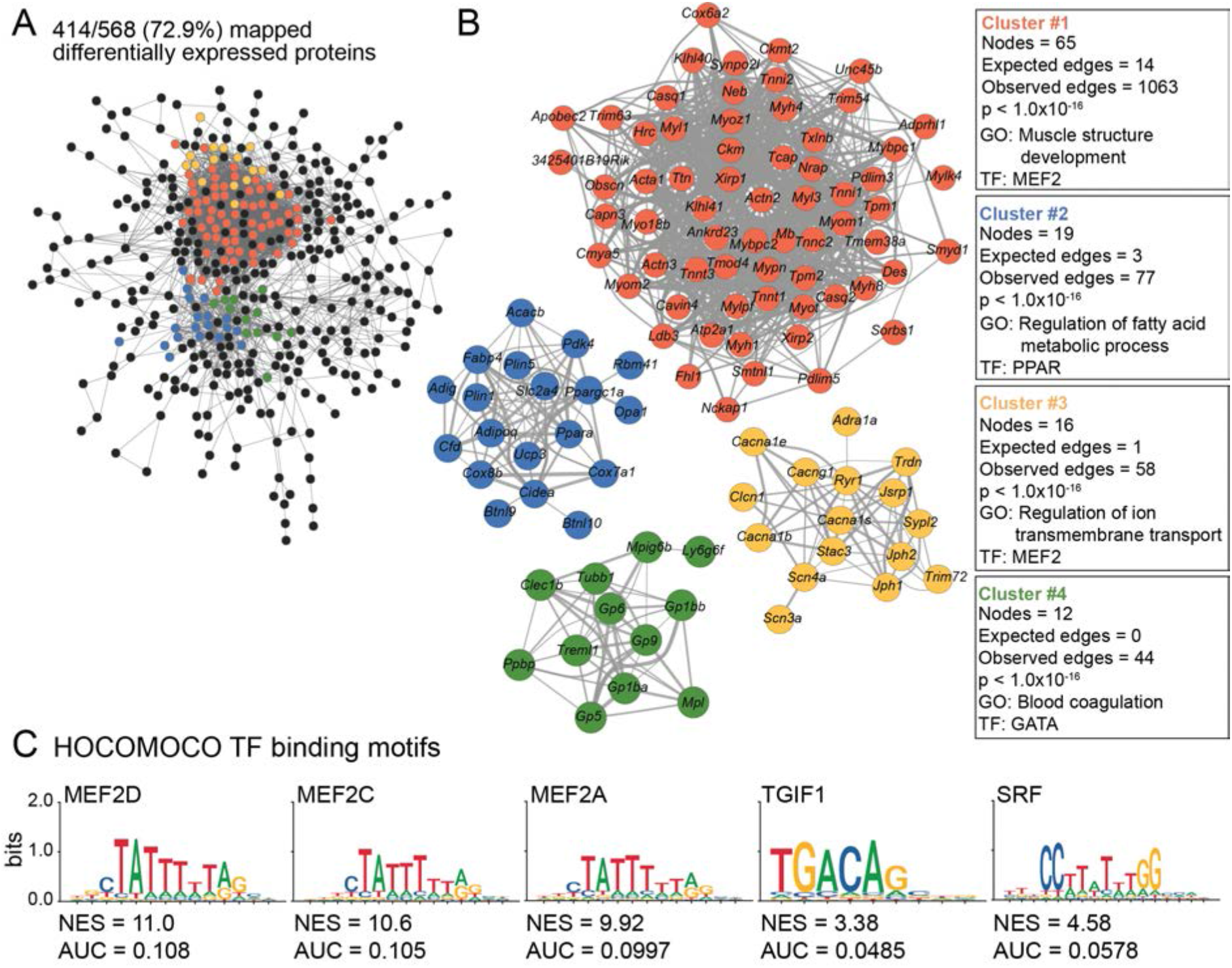
The *Cdh23^ahl^*-associated transcriptional network expands and acquires prominent MEF2-family motif enrichment by P30. **(A)** STRING protein-interaction analysis of DEGs identified between P30 B6N.*Cdh23^ahl^* and B6N.*Cdh23^repair^* cochleae. Of 657 DEGs, 568 mapped to proteins in STRING and 414 formed a large interconnected network. Nodes represent proteins and edges represent known or predicted protein interactions; edge thickness indicates interaction confidence. **(B)** MCL analysis identified four major modules containing >10 proteins within the P30 network. The modules were enriched for biological processes associated with muscle structure development (Cluster 1), regulation of fatty-acid metabolism (Cluster 2), regulation of ion transmembrane transport (Cluster 3), and blood coagulation (Cluster 4). The most highly enriched TF binding-motif family identified by RcisTarget for each cluster is shown. Network visualization was performed using Cytoscape. **(C)** TF binding-motif enrichment across the complete set of 657 P30 DEGs. MEF2-family motifs were among the most highly enriched motifs, together with motifs associated with TGIF1 and SRF. Motif enrichment was assessed within the region spanning 500 bp upstream to 100 bp downstream of the transcription start site using RcisTarget and the HOCOMOCO mouse motif database. NES, normalized enrichment score; AUC, area under the curve. Complete motif-enrichment results are provided in Supplementary Files 4 and 5.

Markov clustering identified four major modules within this network. The largest, Cluster 1, contained 65 proteins and was strongly enriched for genes associated with muscle structure development (adjusted *P* = 6.28 × 10^−36^). Cluster 2 contained 19 proteins and was enriched for regulation of fatty-acid metabolic processes (adjusted *P* = 5.46 × 10^−6^); Cluster 3 contained 16 proteins and was enriched for regulation of ion transmembrane transport (adjusted *P* = 1.55 × 10^−10^); and Cluster 4 contained 12 proteins and was enriched for blood coagulation (adjusted *P* = 1.03 × 10^−11^; Figure 5B; Supplementary File 3). Thus, between P16 and P30, the transcriptional differences associated with *Cdh23^ahl^* expand from a relatively restricted set of immune- and cell-cycle-associated genes to a broader set of interconnected biological programs.

Transcription factor motif analysis using RcisTarget also revealed a distinct regulatory signature at P30. Clusters 1 and 3 were enriched for myocyte enhancer factor 2 (MEF2)-family binding motifs (NES = 13.1 and 6.19, respectively), whereas Cluster 2 was enriched for PPAR-family motifs (NES = 4.91) and Cluster 4 for GATA-family motifs (NES = 9.02; Figure 5B). Across the complete set of 657 P30 DEGs, MEF2-family motifs were among the most highly enriched, including MEF2A (NES = 9.92), MEF2C (NES = 10.6), and MEF2D (NES = 11.0). In total, 246 of the 657 P30 DEGs (37.4%) contained a predicted MEF2-family binding motif within the analyzed transcription-start-site region. TGIF1 (NES = 3.38) and SRF (NES = 4.58) motifs were also enriched (Figure 5C; Supplementary Files 4 and 5).

Together, these analyses reveal an age-dependent evolution of the transcriptional response associated with *Cdh23^ahl^*. Shortly after hearing onset, differential expression is concentrated in interconnected immune- and cell-cycle-associated programs with RUNX- and E2F-family motif enrichment. By P30, the response has expanded into a substantially larger network involving multiple biological processes and is accompanied by prominent enrichment of MEF2-family regulatory motifs. These findings indicate that the molecular consequences associated with *Cdh23^ahl^* are dynamic rather than static, increasing in scale and changing in composition during the interval in which high-frequency auditory dysfunction begins to emerge.

### Age-dependent *Cdh23^ahl^*-associated transcriptional programs are enriched across distinct cochlear cell populations

To define how the transcriptional differences associated with *Cdh23^ahl^* evolve between P16 and P30, we performed a likelihood ratio test (LRT) incorporating the interaction between age and genotype. Unlike the pairwise Wald tests used to identify differentially expressed genes between B6N.*Cdh23^ahl^* and B6N.*Cdh23^repair^* cochleae at each individual age, the LRT identified genes whose expression trajectories differed between genotypes across the two time points. Of the combined 844 DEGs identified at P16 or P30, 395 showed significant age-dependent expression patterns and were assigned to three groups based on similarity of their expression trajectories (Figure 6A; Supplementary File 6). Group 1 contained 98 genes, Group 2 contained 196 genes, and Group 3 contained 101 genes.

**Figure 6.**
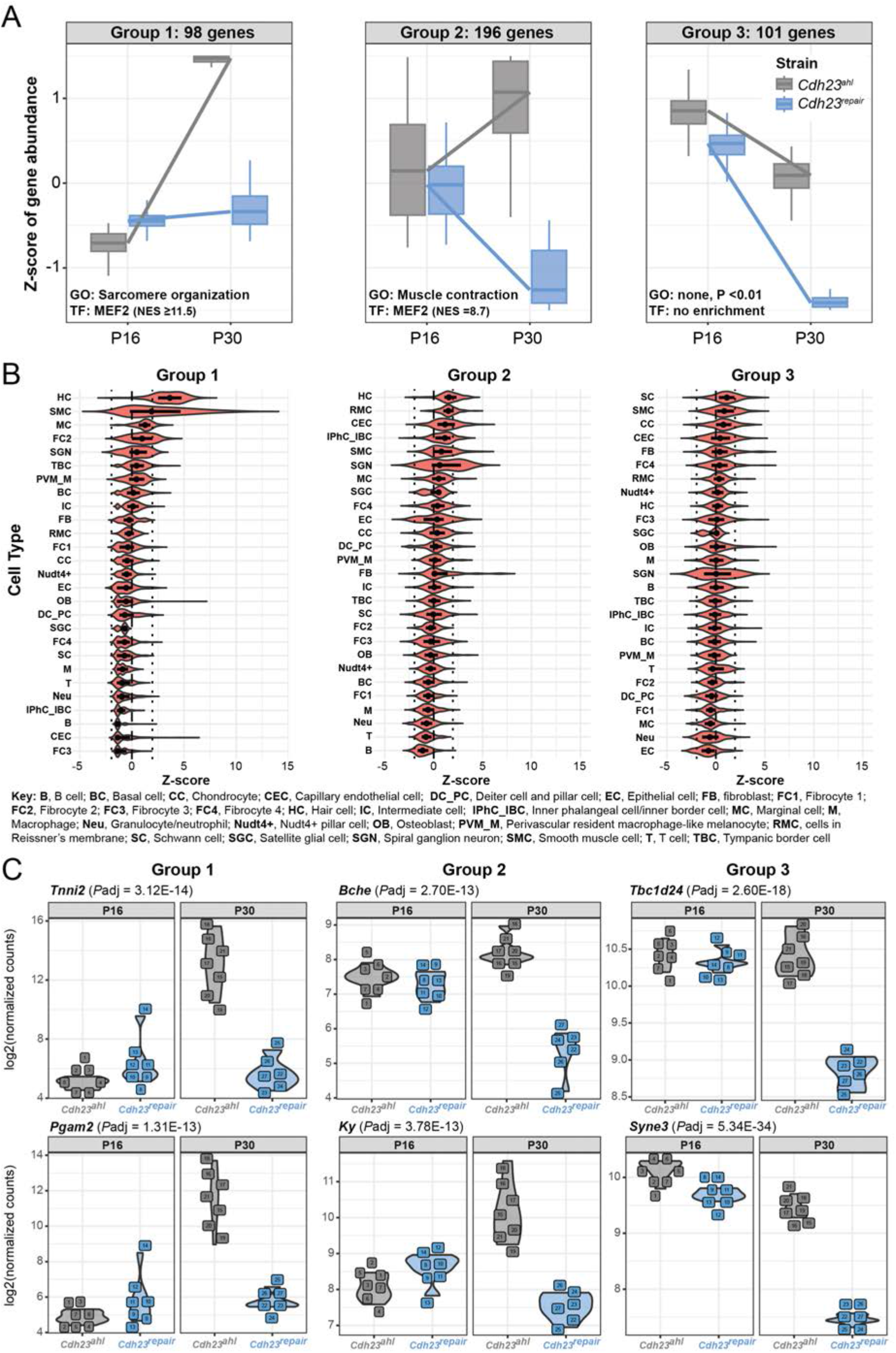
Age-dependent *Cdh23^ahl^*-associated gene programs are represented across distinct cochlear cell populations. **(A)** Likelihood ratio test (LRT) analysis of the combined P16 and P30 DEG sets identified 395 genes with significant genotype-dependent expression trajectories across age. Genes were grouped according to similarity of their temporal expression patterns into Group 1 (98 genes), Group 2 (196 genes), and Group 3 (101 genes). Group 1 was enriched for genes associated with sarcomere organization and Group 2 for genes associated with muscle contraction; no significantly enriched Gene Ontology biological-process term was identified for Group 3 at adjusted *P* <0.01. Groups 1 and 2 also showed enrichment for MEF2-family TF binding motifs. **(B)** Projection of the three gene-expression signatures onto a published single-cell RNA-seq dataset from one-month-old C57BL/6J cochleae using ProjectR. Projection scores >1.96, corresponding to the upper bound of the 95% confidence interval, were considered strong enrichment for the corresponding gene signature. Group 1 showed prominent enrichment in hair cells and smooth muscle cells, with additional enrichment in marginal cells, type 2 fibrocytes, and subsets of spiral ganglion neurons. Group 2 showed enrichment in several populations including hair cells and Reissner’s membrane cells, whereas Group 3 showed enrichment in populations including Schwann cells and smooth muscle cells. Heterogeneous projection scores were observed among spiral ganglion neurons for all three gene groups. **(C)** Normalized read counts for representative genes from Groups 1–3 in individual B6N.*Cdh23^ahl^*and B6N.*Cdh23^repair^* cochlear samples at P16 and P30. Expression among B6N.*Cdh23^ahl^* samples becomes more variable at P30 for representative genes from Groups 1 and 2.

Functional enrichment analysis revealed distinct properties of these expression groups. Group 1 was enriched for genes associated with sarcomere organization (adjusted *P* = 8.94 × 10^−15^), whereas Group 2 was enriched for genes associated with muscle contraction (adjusted *P* = 1.31 × 10^−10^). No Gene Ontology biological-process term reached an adjusted *P* value <0.01 for Group 3. Consistent with the regulatory signatures identified among the P30 DEGs, Groups 1 and 2 showed strong enrichment for MEF2-family transcription factor binding motifs (NES ≥11.5 and 8.7, respectively), whereas Group 3 showed no significant enrichment for a HOCOMOCO transcription factor binding motif.

Due to these analyses being performed on bulk cochlear RNA-seq data, we next sought to determine the cochlear cell populations in which the three gene expression signatures are preferentially represented. We used ProjectR to project each gene group onto a published single-cell RNA-seq dataset generated from one-month-old C57BL/6J cochleae^28^. Projection scores >1.96, corresponding to the upper bound of a 95% confidence interval, were considered evidence of strong enrichment for the respective gene signature.

The Group 1 signature showed enrichment across several cochlear cell populations, most prominently hair cells and smooth muscle cells, with additional enrichment in marginal cells, type 2 fibrocytes, and subsets of spiral ganglion neurons (Figure 6B). Group 2 showed enrichment in several populations, including hair cells and Reissner’s membrane cells, whereas the Group 3 signature was enriched in populations including Schwann cells and smooth muscle cells.

All three gene groups also showed heterogeneous enrichment among spiral ganglion neurons, with high projection scores in some neurons and low scores in others (Figure 6B). This pattern raises the possibility that the transcriptional programs are preferentially represented in specific spiral ganglion neuron populations, although the present analysis does not resolve whether this reflects defined neuronal subtypes.

Examination of normalized read counts for representative genes from each expression group further demonstrated the increasing inter-individual variability observed in the bulk transcriptomic data (Figure 6C). B6N.*Cdh23^repair^* samples showed relatively consistent expression at both ages. B6N.*Cdh23^ahl^*samples were similarly consistent at P16, but showed greater inter-individual variability at P30, particularly among representative genes from Groups 1 and 2.

Together, these analyses indicate that the age-dependent transcriptional programs associated with *Cdh23^ahl^* are not restricted to a single cochlear cell population. Instead, their constituent genes are preferentially represented across multiple cochlear cell types, including sensory hair cells as well as non-sensory and neuronal populations. The enrichment of Groups 1 and 2 in hair cells, together with their strong MEF2-family motif enrichment, provides a potential cellular context for the prominent MEF2-associated regulatory signature that emerges by P30.

## Discussion

The *Cdh23^ahl^* allele has long been recognized as a major contributor to progressive hearing loss in C57BL/6-derived mouse strains^29–31,12^. Our findings demonstrate that its biological consequences begin substantially earlier than previously assumed. By comparing C57BL/6N mice with a co-isogenic strain in which *Cdh23^ahl^* was repaired, we identify physiological and molecular differences shortly after the onset of hearing, before substantial auditory dysfunction is apparent. These early changes are followed by striking inter-individual variability in both cochlear transcriptional state and the onset and severity of high-frequency hearing loss. Thus, *Cdh23^ahl^* should not be considered a physiologically benign background allele in young mice. Rather, our findings support a model in which this hypomorphic allele establishes an altered cochlear state that precedes progressive auditory dysfunction and may influence the phenotypic consequences of additional genetic perturbations.

### The auditory consequences of *Cdh23^ahl^* begin earlier and are more heterogeneous than generally appreciated

Progressive hearing loss in C57BL/6-derived strains carrying *Cdh23^ahl^* is well established and is characterized by an initial loss of high-frequency sensitivity followed by progressive involvement of lower frequencies. However, this phenotype is commonly considered to become apparent at approximately three months of age or later. Our analysis of a large longitudinal ABR dataset, together with measurements extending to 42 kHz, demonstrates that the auditory phenotype begins substantially earlier. By one month of age, B6N.*Cdh23^ahl^* mice already exhibit significant threshold elevations at 36 and 42 kHz, whereas thresholds at lower frequencies remain comparable to, although more variable than, those of B6N.*Cdh23^repair^* mice.

Equally important is the marked inter-individual variability in the auditory phenotype. By three months of age, 32 kHz thresholds among B6N.*Cdh23^ahl^* mice span the range from normal hearing to no detectable response at the maximum stimulus level, whereas thresholds in B6N.*Cdh23^repair^* mice remain comparatively homogeneous. This variability persists as hearing loss progresses before narrowing at later ages as increasing numbers of B6N.*Cdh23^ahl^*mice develop severe hearing impairment. These observations suggest that genetically identical mice carrying *Cdh23^ahl^* do not progress synchronously from normal hearing to hearing loss but instead follow markedly different trajectories.

The source of this variability remains unknown. We found no consistent effect of biological sex and no relationship between high-frequency thresholds and litter or early postnatal cage density. Other environmental, developmental, or stochastic factors may therefore determine when individual *Cdh23^ahl^* cochleae transition toward dysfunction. Regardless of its origin, this variability has practical consequences for experimental design. Studies of genetically modified mice maintained on *Cdh23^ahl^*-containing backgrounds may attribute threshold differences to an experimental mutation when they instead reflect, or are amplified by, variation in the underlying *Cdh23^ahl^*-associated trajectory.

### *Cdh23^ahl^* alters mechanotransduction rather than simply reducing MET current size

The early basal-cochlear vulnerability is particularly interesting considering the essential role of CDH23 in the hair-cell mechanotransduction apparatus. CDH23 forms the upper portion of the stereociliary tip link and is therefore positioned to influence the conversion of mechanical force into MET-channel opening. A simple model for *Cdh23^ahl^*-associated hearing loss would predict that altered CDH23 reduces tip-link function and consequently decreases mechanotransduction. However, while it is reported that *Cdh23^ahl^* mice show decreasing MET current size with age in the apical region (9-12 kHz)^32–34^, our basal measurements do not support such a uniform loss-of-function model.

At P14, shortly after hearing onset and before the high-frequency ABR phenotype measured at one month, maximal MET currents were altered in B6N.*Cdh23^ahl^*IHCs in a tonotopically distinct manner. Currents were smaller in apical, low-frequency, IHCs but, unexpectedly, larger in basal, high-frequency, IHCs compared with B6N.*Cdh23^repair^* controls. In contrast, resting open probability and adaptation kinetics were not detectably altered. Thus, *Cdh23^ahl^* does not abolish or simply reduce MET currents throughout the cochlea. Instead, it alters the magnitude of the transduction current in a manner that differs along the tonotopic axis. This aligns with the fact that *Cdh23^ahl^*does not cause congenital deafness.

The increased maximal MET current in basal IHCs is particularly notable because this is the cochlear region in which auditory dysfunction subsequently emerges first. The present experiments do not establish that it is the observed increased MET current that causes the subsequent high-frequency hearing loss. Nevertheless, our data raise the possibility that altered mechanotransduction contributes to an increased vulnerability towards dysfunction at the base of the cochlea. Increased current could reflect changes in the number or conductance of functional MET channels, tip-link or bundle architecture, or a compensatory response to altered CDH23 function. It could also alter ionic loading or other downstream processes in mechanically active hair cells. Distinguishing among these possibilities will require direct assessment of the underlying MET machinery and longitudinal experiments linking early physiological abnormalities to later hair-cell dysfunction.

### Transcriptomic differences precede substantial hearing loss and become increasingly heterogeneous

The transcriptomic data provide independent evidence that *Cdh23^ahl^*alters cochlear biology early in life. At P16, only two days after the electrophysiological measurements and shortly after hearing onset, B6N.*Cdh23^ahl^* and B6N.*Cdh23^repair^*cochleae already occupy distinct transcriptional states. This finding is important because it demonstrates that the apparent preservation of conventional auditory thresholds early in life does not reflect a biologically normal cochlea. Molecular consequences of the allele are detectable before substantial threshold elevations emerge.

These transcriptional differences increase considerably between P16 and P30. The number of differentially expressed genes increases from 187 at P16 to 657 at P30, with the latter response dominated by increased gene expression. The composition of the response also changes. At P16, differentially expressed genes form networks enriched for immune-system and cell-cycle-associated processes, with corresponding enrichment of RUNX- and E2F-family binding motifs. By P30, the response encompasses a substantially larger interconnected network involving gene sets associated with fatty-acid metabolism, ion transport, and coagulation, together with prominent enrichment of MEF2-family motifs. These observations indicate that the molecular consequences associated with *Cdh23^ahl^* evolve over time rather than representing a fixed transcriptional signature.

The motif-enrichment analyses identify candidate regulatory programs that may contribute to this evolving response, but should not be interpreted as evidence that the corresponding transcription factors directly drive the observed expression changes. The prominent enrichment of MEF2-family motifs at P30 is notable because MEF2 factors have previously been implicated in cochlear development and auditory-neuronal function^35^, and MEF2D has been linked experimentally to regulation of inner-hair-cell ribbon synapses^36,37^. These observations raise the possibility that MEF2-associated transcriptional responses contribute to the evolving cochlear state in *Cdh23^ahl^*mice. However, motif enrichment does not establish MEF2 occupancy or activity, and direct measurements will be required to determine whether MEF2 factors regulate the transcriptional changes identified here.

A particularly striking feature of the P30 data is the increased inter-individual dispersion of the B6N.*Cdh23^ahl^* transcriptomes. Whereas repaired cochleae remain relatively homogeneous, individual *Cdh23^ahl^*cochleae diverge markedly from one another by one month of age. This molecular heterogeneity emerges during the same developmental interval in which auditory thresholds also become increasingly variable. Because transcriptomic and auditory measurements were not obtained from the same animals, we cannot determine whether mice with the most divergent molecular profiles are those with the greatest auditory dysfunction. Nevertheless, the parallel emergence of molecular and functional heterogeneity suggests that variability in the biological response to *Cdh23^ahl^* may contribute to variability in phenotypic expressivity.

Projection of the age-dependent gene-expression signatures onto an independent cochlear single-cell RNA-seq dataset further suggests that these molecular programs are represented across multiple cochlear cell populations rather than being restricted to a single cell type. Hair cells were prominently enriched for several signatures, consistent with the known function of CDH23 and the early MET phenotype, but enrichment was also observed in smooth muscle cells, marginal cells, fibrocytes, Reissner’s membrane cells, Schwann cells, and subsets of spiral ganglion neurons. Because these assignments are based on projection of bulk-derived signatures onto an external single-cell reference dataset, they do not establish which cell types undergo differential expression in B6N.*Cdh23^ahl^* mice. Single-cell or spatial profiling directly comparing *Cdh23^ahl^* and repaired cochleae will be required to resolve the cellular origins and trajectories of these responses.

### An altered baseline state may explain the genetic-modifier effects of *Cdh23^ahl^*

These findings may help explain how *Cdh23^ahl^*: modifies the phenotypic consequences of *Neuroplastin*^19^, *Embigin*^22^*, Sod1*^17^, *Atp2b2*^38^, *Ocm*^20^, and *Wbp2*^21^. Our findings suggest that these interactions could arise from *Cdh23^ahl^*-induced changes in cellular physiology and gene expression before overt hearing loss develops. A second mutation would therefore act within an already altered cellular environment, potentially changing its phenotypic effects. Variation between individuals in the extent of these changes could also contribute to incomplete penetrance or variable expressivity.

This model remains to be tested directly, particularly for phenotypes outside the auditory system, but it provides a potential explanation for why repair of *Cdh23^ahl^* can markedly alter the severity of otherwise identical genetic models. More broadly, it illustrates how a common hypomorphic allele within an inbred genetic background can influence interpretation of experimentally introduced mutations even when the background allele itself produces little overt phenotype at the age being studied.

### Effects associated with *Cdh23^ahl^* may extend beyond the auditory system

Broad phenotyping identified relatively few non-auditory differences between B6N.*Cdh23^ahl^* and B6N.*Cdh23^repair^* mice maintained under standard conditions. The most prominent was lens opacity, which was present in approximately 40% of B6N.*Cdh23^ahl^* mice assessed at 58 weeks, but absent from the repaired cohort. The incomplete penetrance of this phenotype parallels the broader theme of variable expressivity associated with *Cdh23^ahl^* and may be relevant to studies that rely on visual behavioural paradigms in aged C57BL/6-derived mice.

The association between *Cdh23^ahl^* and lens opacity was unexpected. Although *CDH23* is present in the eye, it is primarily expressed in the retina^39^, with severe loss-of-function mutations leading to progressive retinitis pigmentosa (Usher syndrome type 1D) in addition to deafness^8,10,11^. However, there have been reports of an increased risk of cataracts in patients with *CDH23*-associated Usher syndrome^40^, suggesting that *CDH23*-associated disease may affect the lens indirectly. Independent replication and longitudinal ocular characterization will be required to determine the mechanism and specificity of this phenotype.

The relative absence of additional systemic phenotypes under standard housing conditions does not exclude broader modifier effects of *Cdh23^ahl^*. Indeed, previous genetic-interaction studies suggest that some consequences of the allele may become apparent primarily when the biological system is challenged by another genetic or environmental perturbation^22^. Testing the repaired and *ahl* backgrounds under defined metabolic, inflammatory, environmental, or genetic challenges could help establish the extent to which *Cdh23^ahl^* modifies phenotypic resilience beyond the cochlea.

### Implications and limitations

Several limitations are important when interpreting these findings. First, although the auditory and transcriptomic datasets independently demonstrate substantial inter-individual variability, they were generated from different animals and therefore cannot establish a direct relationship between molecular state and hearing phenotype. Second, bulk cochlear RNA sequencing cannot distinguish changes in gene expression within individual cell types from changes in cellular composition. The single-cell projection analysis provides hypotheses regarding cellular localization, but does not resolve this limitation. Third, transcription factor motif enrichment identifies candidate regulatory relationships, but does not demonstrate transcription factor binding or activity. Finally, the mechanisms responsible for the opposite effects of *Cdh23^ahl^* on apical and basal IHC MET-current amplitude remain unknown.

These limitations point directly to future experiments. Paired longitudinal auditory phenotyping followed by single-cell or spatial molecular profiling could determine whether molecular heterogeneity predicts subsequent auditory decline. Direct interrogation of the MET apparatus in apical and basal hair cells could establish why *Cdh23^ahl^*produces tonotopically divergent current amplitudes. Finally, comparison of genetically modified lines on *Cdh23^ahl^* and repaired backgrounds could test whether the early molecular state identified here predicts the direction or magnitude of genetic interactions.

In summary, our findings redefine the biological window during which *Cdh23^ahl^* influences cochlear function. Rather than acting only as a determinant of later-onset hearing loss, *Cdh23^ahl^* alters hair-cell physiology and cochlear gene expression shortly after hearing onset, followed by increasing molecular and auditory phenotypic heterogeneity. These findings provide a potential framework for understanding the variable expressivity and genetic-modifier effects associated with this common hypomorphic allele. Given the widespread use of C57BL/6-derived and other mouse strains carrying *Cdh23^ahl^*, accounting for these early and heterogeneous effects will be important for the design and interpretation of studies of auditory function and genetically modified mouse models.

## Methods

### Mouse Models

All animal studies were licensed by the Home Office under the Animals (Scientific Procedures) Act 1986, United Kingdom, and additionally approved by Institutional Ethical Review Committees at both the Mary Lyon Centre, and the University of Sheffield. Mice were housed under a 12-hour light/12-hour dark cycle and allowed access to food and water ad libitum. All conducted experiments complied with the journal’s ethics policies.

The C57BL/6NTac.*Cdh23^repair^* mouse (B6N.*Cdh23^repair^*, RRID:MGI:5749861) was produced at the Mary Lyon Centre at MRC Harwell using CRISPR/Cas9-mediated homology-directed repair to correct the *Cdh23* allele to “wild type” (*Cdh23^c.753A>G^*)^14^. This line was compared with the standard C57BL/6NTac background (B6N.*Cdh23^ahl^*, RRID:IMSR_TAC:b6). To ensure that these lines remained co-isogenic, the C57BL/6NTac.*Cdh23^repair^* strain is bred for no more than four generations before crossing back onto the standard C57BL/6NTac background. Both male and female mice were used in these studies.

### Standardised Phenotyping Pipeline

Open Field, Grip Strength, Combined SHIRPA and Dysmorphology, Echocardiogram (Echo-CG), Electrocardiogram (ECG), Intraperitoneal glucose tolerance test (IPGTT), Body Composition (DEXA lean/fat), Eye Morphology, and collection of tissues and blood for pathology, haematology, and clinical chemistry were performed the Mary Lyon Centre at MRC Harwell. Full procedural details are available via the International Mouse Phenotyping Resource of Standardised Screens (IMPReSS) under the Harwell late adult pipeline (HRWLLA_001)^41,42^.

### Auditory Brainstem Response (ABR) Recordings

Mice were anesthetized using intraperitoneal injection of ketamine (100 mg/Kg body weight, Fort Dodge Animal Health, USA) and xylazine (10 mg/Kg, Rompun, Bayer HealthCare LLC, USA). At the end of the ABR recordings, mice were either culled by cervical dislocation or recovered from anaesthesia with intraperitoneal injection of atipamezole (1 mg/Kg, Antisedan, Orion Corporation, Finland). Following the onset of anaesthesia and the loss of the retraction reflex with a toe pinch, mice were placed in a soundproof chamber (MAC-3 acoustic chamber, IAC Acoustic, UK). Male and female mice were placed on a heated mat (37°C) with the animal’s pinna being positioned at a distance of 10 cm from the speaker. Two subdermal electrodes were placed under the skin behind the pinna of each ear for the reference and ground electrode, and one electrode was placed between the two pinna on the vertex of the cranium as an active electrode. ABR responses were collected, amplified, and averaged using the TDT System 3 (Tucker Davies Technology, Alachua, FL, USA) in conjunction with BioSig RZ software. Stimuli were delivered in the form of a 0.1 ms broadband click, or as a single frequency tone at 3, 6, 8, 12, 16, 18, 24, 30, 36, and/or 42 kHz (5 ms duration, 1 ms rise and fall). All stimuli were presented in 5 dB falling steps from 95 dB SPL, and responses were averaged over 300-1000 repeats.

### Analysis of auditory phenotypes

Auditory brainstem response (ABR) thresholds were compiled from experiments performed between 2016 and 2023 in which B6N.*Cdh23^ahl^* and B6N.*Cdh23^repair^* mice were assessed using 8-, 16-, and 32-kHz pure-tone stimuli at one or more ages between 1 and 18 months. Because a subset of animals underwent repeated measurements, individual mouse identity was retained throughout the analysis.

Differences in threshold distributions between genotypes at individual ages and frequencies were assessed using non-parametric comparisons with correction for multiple testing, as specified in the figure legends. To test whether *Cdh23^ahl^* was associated with increased inter-individual variability, dispersion was analyzed separately by age and stimulus frequency using absolute deviations from the genotype-specific median using a Brown-Forsythe test. Where appropriate, experimental cohort/year was evaluated as a potential source of variation because measurements were pooled across studies performed between 2016 and 2023.

### Tissue preparation for ex vivo patch-clamp recordings

Cochleae were dissected out in extracellular solution composed of (in mM): 135 NaCl, 5.8 KCl, 1.3 CaCl2, 0.9 MgCl2, 0.7 NaH2PO4, 5.6 D-glucose, 10 HEPES-NaOH. Sodium pyruvate (2 mM), amino acids and vitamins were added from concentrates (Thermo Fisher Scientific, UK). The pH was adjusted to 7.48 with 1M NaOH, and the final osmolality of the solution was measured (∼306 mmol/kg). The dissected apical coil was then transferred to a microscope chamber and immobilised using a nylon mesh. The chamber was perfused from a peristaltic pump (Cole-Palmer, UK) and mounted on the stage of an upright microscope (Olympus BX51, Japan). The individual stereocilia rows of the hair cells was identified using a 60x water immersion objective and a 15x eyepiece.

### Whole-cell electrophysiology

Patch clamp recordings were performed at room temperature (20-24°C) using an Optopatch amplifier (Cairn Research Ltd, UK). Patch pipettes were pulled from soda glass capillaries and had a typical resistance in extracellular solution of 2-3 MΩ. The patch pipette intracellular solution contained (in mM): 131 CsCl, 3 MgCl2, 1 EGTA-KOH, 5 Na2ATP, 5 HEPES-CsOH, 10 Na-phosphocreatine (pH was adjusted with 1M CsOH to 7.28, ∼290 mmol/kg). Data acquisition was controlled by pClamp software using Digidata 1550 (Molecular Devices, USA). To reduce the electrode capacitance, patch electrodes were coated with surf wax (Mr. Zoggs SexWax, USA). Recordings were low-pass filtered at 2.5 kHz (8-pole Bessel), sampled at 5 kHz and stored on a computer for off-line analysis (Clampfit, Molecular Devices and Origin 2020, OriginLab). For voltage-clamp, membrane potentials were corrected off-line for the liquid junction potential (LJP) of −4 mV, which was measured between electrode and bath solutions. Holding currents were plotted as zero current to allow a better comparison between recordings. Voltage clamp protocols are referred to a holding potential of −84 mV.

For mechanoelectrical transducer (MET) current recordings, the hair bundles of hair cells were displaced using a fluid jet from a pipette driven by a 25 mm diameter piezoelectric disc. The pipette was pulled from borosilicate glass to a final overall length of 5.3-5.5 cm. The fluid jet pipette tip had a diameter of 8-10 µm and was positioned near the hair bundles to elicit a maximal MET current. Mechanical stimuli were applied as 50 Hz sinusoids (filtered at 1 kHz, 8-pole Bessel). Prior to the positioning of the fluid jet by the hair bundles, any steady-state pressure was removed by monitoring the movement of debris in front of the pipette. The use of the fluid-jet allows for the efficient displacement of the hair bundles in both the excitatory and inhibitory directions, which is essential to perform reliable measurements of the resting open probability of the MET channels. The number of cells recorded is indicated in the figures, and each cell was recorded from a different animal. Apical-coil hair cells were recorded from 9-12 kHz region and basal-coil hair cells were recorded from 40-45 kHz region.

### Cochlear RNA Isolation

From B6N.*Cdh23^ahl^* and B6N.*Cdh23^repair^*mice at P16 and P30, whole cochlear tissues were dissected and the vestibular system removed. Two cochleae (from one mouse) were combined into one sample, immediately submerged in TRIzol™ Reagent (Invitrogen, # 15596026), and snap-frozen on dry ice. Homogenization was performed within Precellys Lysing bead tubes (CK14, 2ml) agitated for 3×20s at 6000RPM with a 30s pause using the Precellys® Evolution Touch tissue homogenizer with the Cryolysis Evolution cooling unit set to 4°C. Following homogenisation, RNA was extracted using the Direct-Zol RNA mini prep kit (ZymoResearch, R2052) with a modified protocol based on methods previously described^43^. Briefly, following homogenisation RNA is separated using chloroform (150µl chloroform for every 750µl of Trizol). Following centrifugation, the aqueous phase is separated and precipitated with equal volumes of EtOH before washing in the RNA mini prep columns. DNAse treatment was performed by incubating the extracted RNA samples with DNAse I (Invitrogen # 10636153). Sample concentrations, purity, and RNA integrity was measured on the Agilent 2100 Bioanalyzer using an Agilent RNA 6000 Nano Kit. RNA quality was assessed before library preparation, and all samples used for sequencing had RNA integrity numbers (RINs) ≥8.5.

### Bulk RNA sequencing

At P16, RNA-seq was performed on seven B6N.*Cdh23^ahl^* cochlear samples (4 male, 3 female) and seven B6N.*Cdh23^repair^*samples (3 male, 4 female). At P30, seven B6N.*Cdh23^ahl^* samples (3 male, 4 female) and six B6N.*Cdh23^repair^* samples (4 male, 2 female) were analyzed. RNA libraries were prepared using the KAPA mRNA HyperPrep Kit (Roche, Switzerland) and were sequenced on a Novaseq 6000 (Illumina, California, USA) using a 100 bp paired end read configuration at the UCL Genomics facility. RNA sequencing data was processed through nf-core/rnaseq (version 3.14.0)^44,45^. Briefly, the reads were trimmed using Trimgalore (version 0.6.7), mapped to GRCm39 using STAR aligner (version 2.7.9a) and quantified with Salmon (version 1.10.1) using Ensembl release 112 as a reference transcript database. All other parameters were used as default. The data discussed in this publication have been deposited in NCBI’s Gene Expression Omnibus and are accessible through GEO Series accession number GSE335571 (https://www.ncbi.nlm.nih.gov/geo/query/acc.cgi?acc=GSE335571). The data are also available on the um-gEAR: Gene Expression Analysis Resource portal^46^ (https://umgear.org/p?l=bl6repair).

### Analysis of RNA-seq data

The BAM files generated by STAR were merged, sorted and indexed by age and strain with SAMtools^47^ for visualization on Integrative Genomics Viewer (IGV)^48^. The *Cdh23* gene was inspected to assess splicing events involving exons 8, 9 and 10 of *Cdh23* transcript ENSMUST00000105461. To rule out mapping error, junctions with at least 10 supporting reads were considered. Gviz^49^ was used to create the final sashimi and coverage plots. Differential expression analysis was performed with DESeq2 (version 1.44.0)^50^, applying a Wald test, for pairwise comparisons of B6N.*Cdh23^repair^* and B6N.*Cdh23^ahl^ at P16 or P30*. Genes with fewer than 10 reads in at least one sample were removed from the analysis. At each age, differential expression between B6N.Cdh23*^ahl^*and B6N.Cdh23*^repair^* cochleae was assessed using the following design formula ∼ sex + genotype. A gene was considered differentially expressed if it presented an adjusted p < 0.05 and absolute logFC ≥ 1. Principal component analysis was performed using a variance stabilizing transformation (vst) on count data values. PCA was performed separately for each age group, using the 500 most variable genes in that analysis.

### Analysis of transcriptomic heterogeneity

Variance-stabilised gene-expression values obtained using DESeq2^50^ were used to identify the 500 most variable genes across B6N.Cdh23*^ahl^* and B6N.Cdh23*^repair^* samples separately at P16 and P30.To quantify inter-individual transcriptomic variability at P16 and P30, a permutational analysis of multivariate dispersion (PERMDISP) was implemented using the betadisper() function in the R package vegan (version 2.7-3)^51^. Euclidean distances between samples were calculated from the expression profiles of these genes, and multivariate dispersion was quantified as the distance of each sample to its genotype-specific centroid. Differences in dispersion between genotypes were assessed using a permutation test with 10,000 permutations. Cohen’s d was calculated using the sample-level distances to group centroids to quantify the standardised difference in dispersion between genotypes.

### Protein-interaction, functional-enrichment, and transcription-factor motif analyses

Protein–protein interaction networks were generated from differentially expressed genes on Cytoscape (version 3.10.3) using the STRING plugin (version 2.1.1)^52,53^. Genes that could be mapped to STRING proteins were used to construct interaction networks using a minimum interaction-confidence score of 0.4. Network enrichment was evaluated by comparing the observed number of interactions with that expected for randomly selected protein sets of equivalent size.

Modules within the STRING networks were identified using the Markov Cluster Algorithm (MCL) with an inflation parameter of 2.5. Clusters containing >10 proteins were analyzed for Gene Ontology biological-process enrichment using StringApp on Cytoscape STRING. Multiple-testing correction was performed using the Benjamini–Hochberg procedure and terms with adjusted *P* <0.01 were considered significantly enriched.

Transcription-factor binding-motif enrichment was assessed using RcisTarget (version 1.24.0)^54^. Motif rankings were based on the HOCOMOCO mouse transcription-factor motif database. For each gene, the region from 500 bp upstream to 100 bp downstream of the transcription start site was examined for enriched motifs. Analyses were performed for the complete P16 and P30 DEG sets and independently for major STRING/MCL clusters and LRT-derived expression groups. Motif enrichment was summarized using the normalized enrichment score (NES).

### Analysis of genotype-dependent expression trajectories

To identify genes whose expression trajectories differed between B6N.*Cdh23^ahl^* and B6N.*Cdh23^repair^* cochleae between P16 and P30, a likelihood ratio test (LRT) was performed using DESeq2. The full model included ∼ sex + age + strain + age:strain, and was compared with a reduced model containing ∼ sex + age + strain. Genes with a Benjamini–Hochberg-adjusted *P* value <0.05 were considered to show significant genotype-dependent temporal expression patterns.

Among genes identified as differentially expressed between genotypes at P16 or P30, 395 met the LRT significance criterion. rlog transformed normalized counts of significant genes from the LRT further were filtered by the DEG lists identified using the Wald test. These genes were grouped according to similarity in their expression trajectories using DEGreport::degpatterns^55^, yielding three groups containing 98, 196, and 101 genes, respectively. Functional enrichment and transcription-factor motif analyses were subsequently performed independently for each group.

### Projection of transcriptional signatures onto cochlear single-cell RNA-seq data

To identify cochlear cell populations in which the age-dependent gene-expression signatures were preferentially represented, the three LRT-derived gene groups were projected onto a published single-cell RNA-seq dataset from one-month-old C57BL/6J cochleae^28^ using ProjectR (version 1.26.0)^56^.

The reference single-cell dataset was obtained from the gEAR portal [https://umgear.org/p?s=728a05e2] Quality control, filtering, normalisation, dimensionality reduction, UMAP embedding and cell-type annotation were performed by the original authors^28^. Each LRT-derived gene signature was projected independently onto the reference single-cell expression matrix using ProjectR, generating a projection score for each cell.

Projection scores >1.96 (standard deviations above the dataset mean) were considered strong enrichment for the corresponding gene signature, which corresponds to the upper 2.5% tail of a standard normal distribution (two-tailed α = 0.05). This was used as a descriptive cut-off rather than a formal test of statistical significance. Projection scores were visualized according to the cell-type annotations reported in the original single-cell dataset. This analysis was used to identify cell populations preferentially expressing genes within each bulk-derived signature and was not used to infer genotype-dependent differential expression within individual cell types.

## Acknowledgements

The authors would like to thank the Phenotyping Team (Mary Lyon Centre, International Facility for Mouse Genetics, at MRC Harwell) for generating the aged phenotyping data.

## Author contributions

Conceptualization S.N., J.J., and M.R.B.,

methodology. S.N., M.F., M.S., S.W., R.H., J.J., M.R.B.,

investigation S.N., M.F., P.G., C.A., J.J.

data analysis S.N., M.F., C.S., P.G., C.A., W.S., J.J., M.R.B.,

writing - original draft S.N., M.F., J.J. and M.R.B.,

writing - review & editing All authors

funding acquisition, S.W., R.H., J.J., and M.R.B.,

supervision, S.W., R.H., and M.R.B.

## Declaration of generative AI technologies

During the preparation of this work no generative AI technologies were used.

## Supplementary

**Supplementary table 1.** descriptive statistics and comparison of inter-animal variability of auditory brainstem response measures at 8, 16, and 32 kHz across age 1–18 months in the B6N.*Cdh23^ahl^* and B6N.*Cdh23^repair^* mice. Data are presented as mean, variance, median, and median absolute deviation (MAD) for each age and frequency. Differences in threshold variance between genotypes were assessed using the median-centred Brown–Forsythe test. Statistically significant differences are indicated by P < 0.05. AHL, B6N.Cdh23*ahl*; Repair B6N.Cdh23*repair*; MAD, median absolute deviation; F, F-statistic.

|  |  | 8Khz |  |  |  |  |  | 16kHz |  |  |  |  |  | 32kHz |  |  |  |  |  |
|---|---|---|---|---|---|---|---|---|---|---|---|---|---|---|---|---|---|---|---|
| age (months) | strain | mean | variance | median | MAD | F | P | mean | variance | median | MAD | F | P | mean | variance | median | MAD | F | P |
| **1** | ahl | 28.3 | 54.7 | 30 | 5 | 8.04 | 6.39E-03 | 20.8 | 37.7 | 20 | 5 | 10.25 | 2.24E-03 | 35.5 | 245.8 | 30 | 5 | 12.03 | 9.20E-04 |
|  | repair | 23.7 | 32.3 | 25 | 5 |  |  | 16.5 | 21.4 | 15 | 5 |  |  | 25.9 | 56.0 | 25 | 5 |  |  |
| **3** | ahl | 26.0 | 40.5 | 25 | 5 | 0.01 | 9.40E-01 | 23.0 | 33.9 | 20 | 5 | 19.24 | 2.08E-05 | 52.9 | 297.5 | 55 | 10 | 212.70 | 4.15E-31 |
|  | repair | 25.9 | 30.4 | 25 | 5 |  |  | 19.5 | 20.9 | 20 | 5 |  |  | 27.0 | 38.1 | 25 | 5 |  |  |
| **6** | ahl | 32.7 | 77.3 | 30 | 5 | 14.57 | 1.85E-04 | 30.7 | 116.5 | 30 | 10 | 47.32 | 1.23E-10 | 70.2 | 170.0 | 70 | 10 | 841.54 | 3.24E-63 |
|  | repair | 28.3 | 47.1 | 30 | 5 |  |  | 22.0 | 37.2 | 20 | 5 |  |  | 28.1 | 37.1 | 25 | 5 |  |  |
| **9** | ahl | 39.7 | 111.7 | 40 | 5 | 29.82 | 2.19E-07 | 41.8 | 264.8 | 40 | 10 | 103.09 | 1.67E-16 | 74.5 | 162.6 | 75 | 10 | 819.81 | 7.88E-48 |
|  | repair | 31.3 | 66.8 | 30 | 5 |  |  | 21.3 | 31.4 | 20 | 5 |  |  | 28.8 | 25.1 | 30 | 5 |  |  |
| **12** | ahl | 49.5 | 100.3 | 50 | 10 | 45.53 | 8.92E-10 | 54.9 | 216.2 | 55 | 15 | 202.44 | 1.02E-21 | 68.8 | 169.5 | 70 | 10 | 436.96 | 5.59E-30 |
|  | repair | 37.5 | 79.7 | 35 | 5 |  |  | 23.9 | 37.5 | 25 | 5 |  |  | 28.9 | 23.7 | 30 | 5 |  |  |
| **15** | ahl | 66.7 | 199.0 | 65 | 10 | 56.24 | 1.24E-11 | 67.8 | 199.5 | 70 | 10 | 408.36 | 2.93E-38 | 76.3 | 75.8 | 75 | 5 | 933.80 | 1.64E-59 |
|  | repair | 50.6 | 97.2 | 50 | 5 |  |  | 27.6 | 59.4 | 25 | 5 |  |  | 31.6 | 59.6 | 30 | 5 |  |  |
| **18** | ahl | 76.1 | 133.3 | 75 | 10 | 17.84 | 3.55E-04 | 74.1 | 194.3 | 80 | 10 | 69.65 | 5.83E-08 | 78.3 | 82.7 | 80 | 5 | 231.46 | 3.91E-12 |
|  | repair | 51.9 | 79.6 | 51.25 | 3.75 |  |  | 32.3 | 53.9 | 30 | 5 |  |  | 36.6 | 57.5 | 35 | 5 |  |  |

**Supplementary Figure 1.**
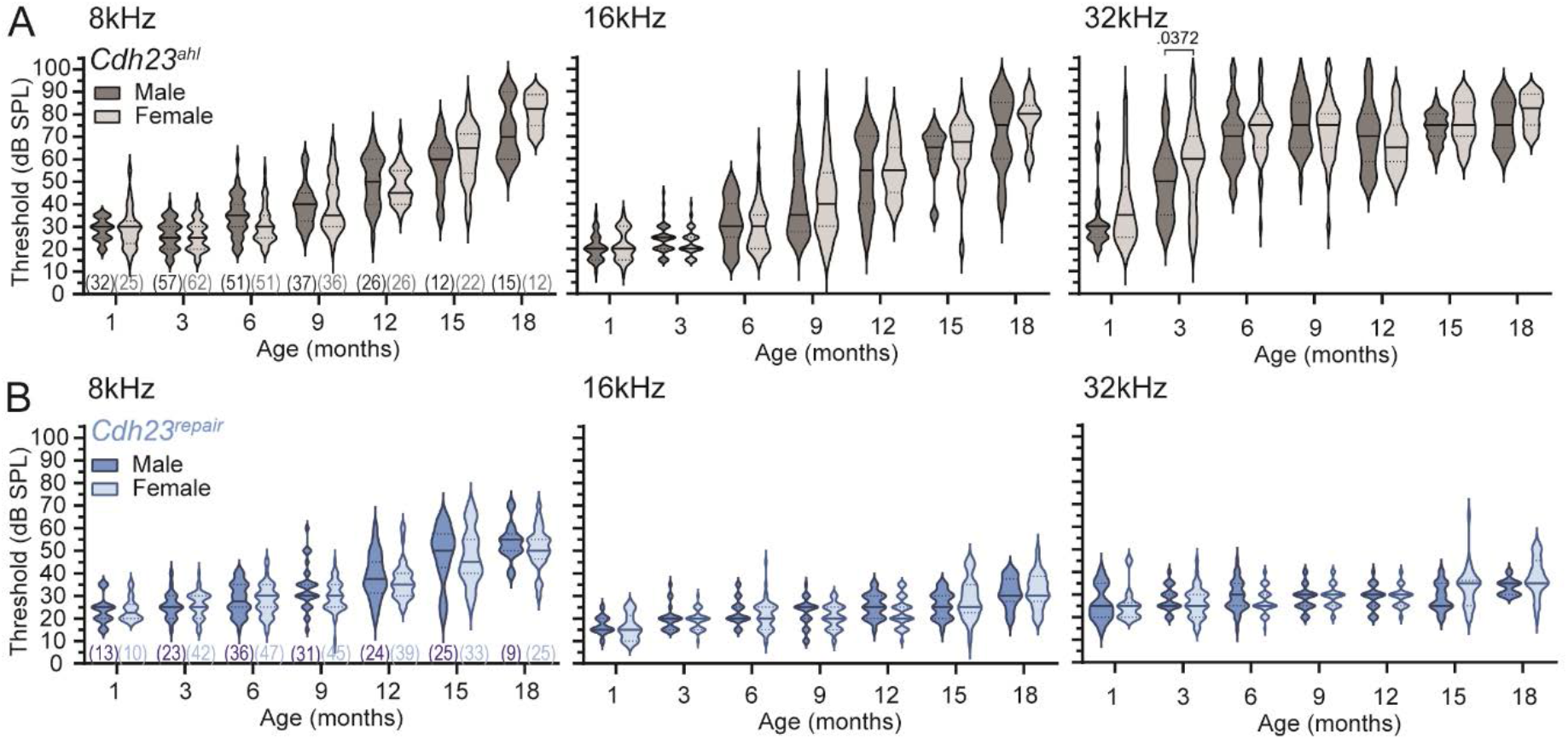
Comparison of male and female ABR measurements (A) B6N.*Cdh23^ahl^* and (B) B6N.*Cdh23^repair^* across all tested timepoints. Violin plots of ABR thresholds recorded for low-(8kHz), mid-(16kHz), and high-frequency stimuli (32kHz). These data were compiled from multiple cohorts recorded within our laboratory between 2016 – 2023. Statistical comparisons in C-D were made using the non-parametric Mann-Whitney test with Holm-Sidak correction for multiple comparisons. P-values for each comparison shown above each plot, and number of mice included shown in brackets below 8kHz plot. No significant differences were observed between sexes in B6N.*Cdh23^repair^* mice. For the B6N.*Cdh23^ahl^* just one difference was observed between males and females 32kHz at three month of age.

**Supplementary Figure 2.**
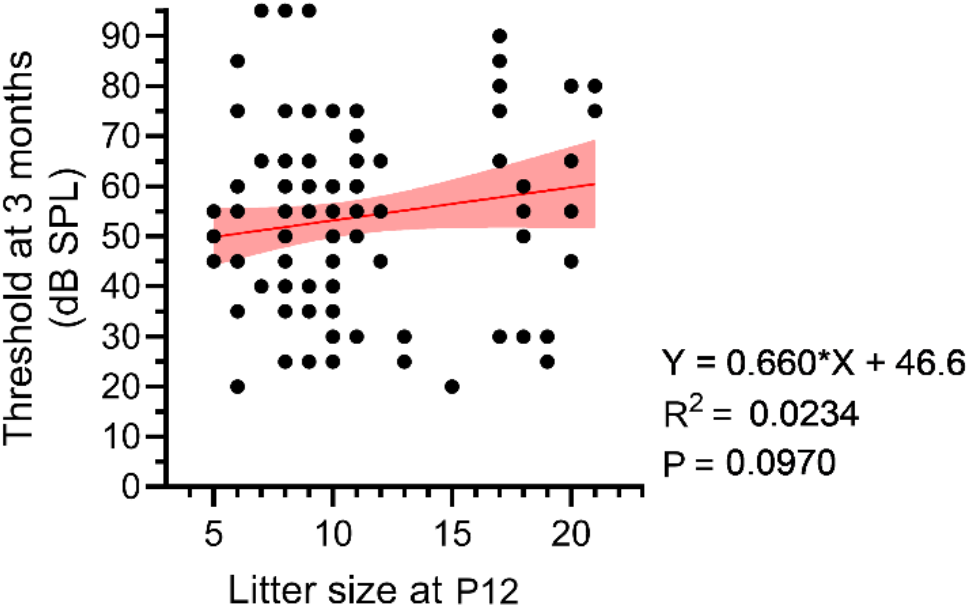
B6N.*Cdh23^ahl^* variation in ABR thresholds at 3-months of age is not associated with litter size. High-frequency (32kHz) auditory brainstem response (ABR) thresholds recorded from B6N.*Cdh23^ahl^* mice at 3-months of age plotted against the litter size they were born in to. In some cases, breeding cages were set up as trios (i.e. two breeding females present) which produced litters around the same time or pups died in the early postnatal period. For this reason, litter size was recorded as the number of pups present in breeding cages between P12 and weaning. There is no correlation between ABR threshold and litter size.

**Supplementary Figure 3.**
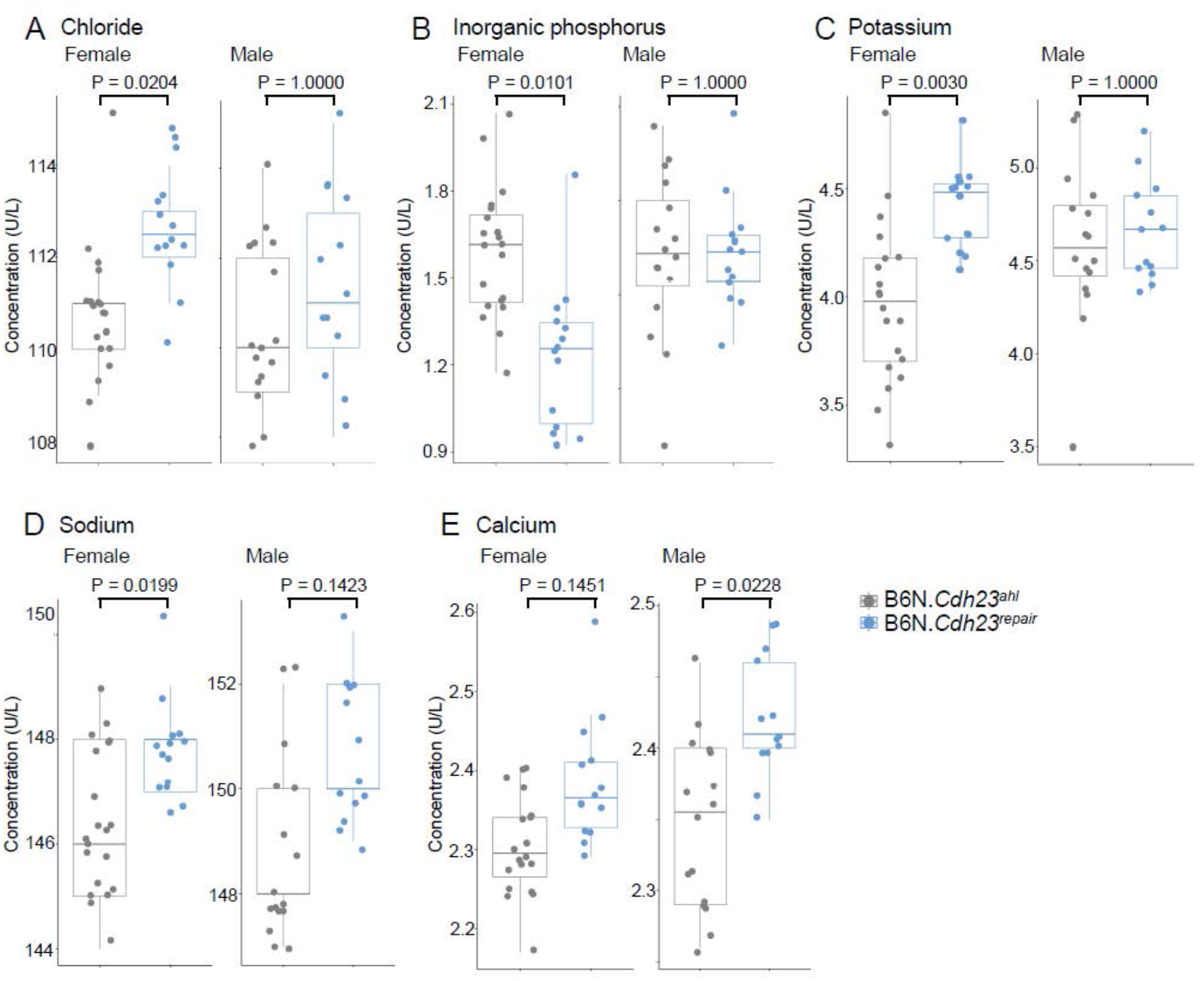
Significant differences in blood clinical chemistry analyses between B6N.*Cdh23^ahl^* and B6N.*Cdh23^repair^* mice collected at 59.5 weeks of age. Of the 21 parameters measured from the collected bloods (see supplementary file 1), significant differences were found in (A) Chloride (females), (B) Inorganic Phosphorus (females), (C) Potassium (females), (D) Sodium (females), and (E) Calcium (males). Comparisons were made using a two-sided t-test with Bonferroni correction for multiple comparisons

**Supplementary Figure 4.**
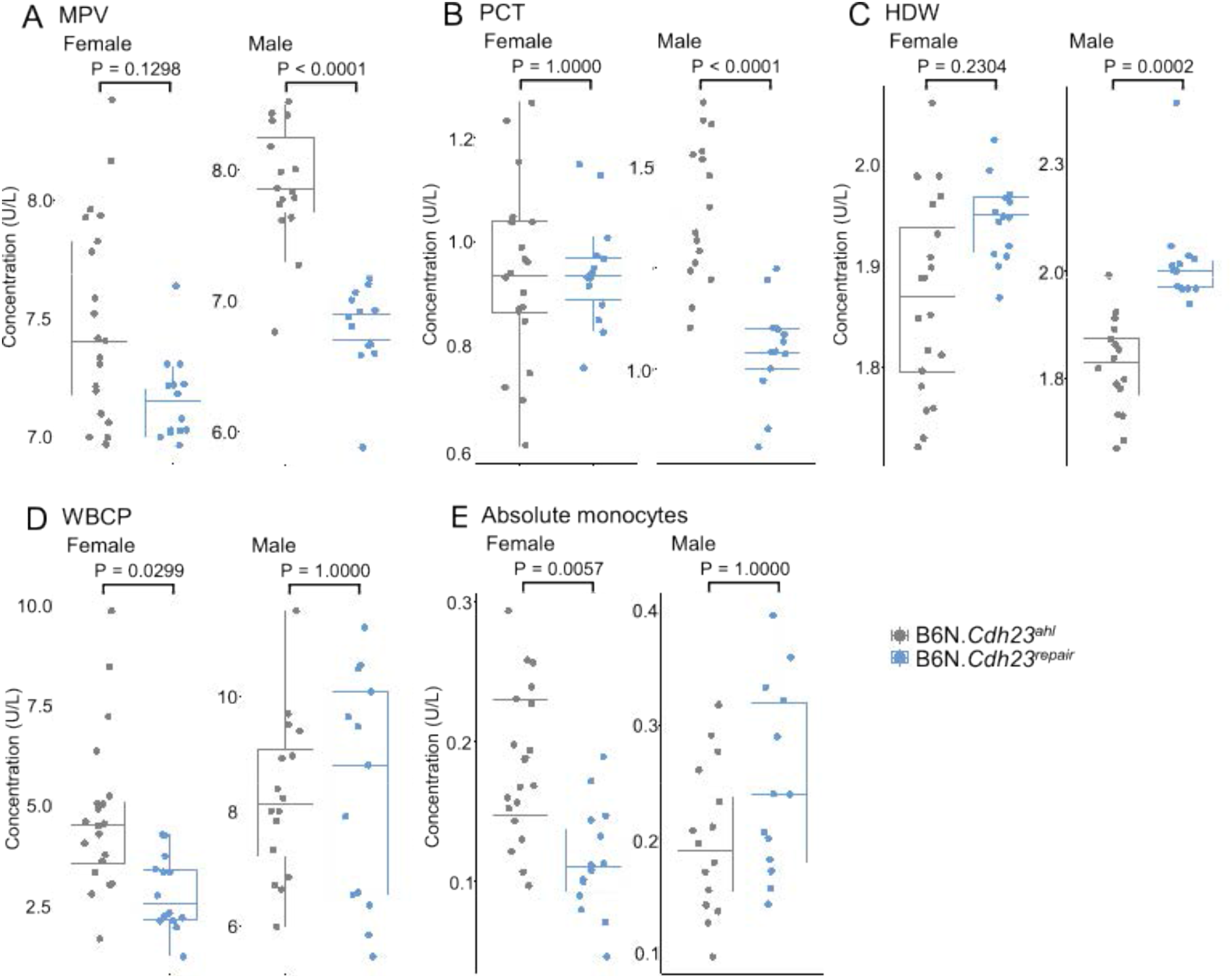
Significant differences in Haematology analyses between between B6N.*Cdh23^ahl^* and B6N.*Cdh23^repair^* mice collected at 59.5 weeks of age. Of the 28 parameters measured from the collected bloods (see supplementary file 1), significant differences were observed in: (A) Mean platelet volumes (MPV, males); (B) Plateletcrit (PTC, males) - a measure of the percentage of blood volume occupied by platelets; (C) Hemoglobin Distribution Width (HDW, males) – a measure of variation in hemoglobin concentration within red blood cells; (D) White Blood Cell count (Peroxidase method, WBCP, females); and (E) absolute monocyte count (females). Comparisons were made using a two-sided t-test with Bonferroni correction for multiple comparisons

